# A modelling framework for the prediction of the herd-level probability of infection from longitudinal data

**DOI:** 10.1101/2020.07.10.197426

**Authors:** Aurélien Madouasse, Mathilde Mercat, Annika van Roon, David Graham, Maria Guelbenzu, Inge Santman Berends, Gerdien van Schaik, Mirjam Nielen, Jenny Frössling, Estelle Ågren, Roger W. Humphry, Jude Eze, George J. Gunn, Madeleine K. Henry, Jörn Gethmann, Simon J. More, Nils Toft, Christine Fourichon

**Author notes:** **Cite as:** Madouasse A, Mercat M, van Roon A, Graham D, Guelbenzu M, Santman Berends I, van Schaik G, Nielen M, Frössling J, Ågren E, Humphry RW, Eze J, Gunn GJ, Henry MK, Gethmann J, More SJ, Toft N, Fourichon C (2020) A modelling framework for the prediction of the herd-level probability of infection from longitudinal data. bioRxiv, 2020.07.10.197426, ver. 6 peer-reviewed and recommended by PCI Animal Science. https://doi.org/10.1101/2020.07.10.197426.

## Abstract

The collective control programmes (**CP**s) that exist for many infectious diseases of farm animals rely on the application of diagnostic testing at regular time intervals for the identification of infected animals or herds. The diversity of these CPs complicates the trade of animals between regions or countries because the definition of freedom from infection differs from one CP to another. In this paper, we describe a statistical model for the prediction of herd-level probabilities of infection from longitudinal data collected as part of CPs against infectious diseases of cattle. The model was applied to data collected as part of a CP against bovine viral diarrhoea virus (**BVDV**) infection in Loire-Atlantique, France. The model represents infection as a herd latent status with a monthly dynamics. This latent status determines test results through test sensitivity and test specificity. The probability of becoming status positive between consecutive months is modelled as a function of risk factors (when available) using logistic regression. Modelling is performed in a Bayesian framework, using either Stan or JAGS. Prior distributions need to be provided for the sensitivities and specificities of the different tests used, for the probability of remaining status positive between months as well as for the probability of becoming positive between months. When risk factors are available, prior distributions need to be provided for the coefficients of the logistic regression, replacing the prior for the probability of becoming positive. From these prior distributions and from the longitudinal data, the model returns posterior probability distributions for being status positive for all herds on the current month. Data from the previous months are used for parameter estimation. The impact of using different prior distributions and model implementations on parameter estimation was evaluated. The main advantage of this model is its ability to predict a probability of being status positive in a month from inputs that can vary in terms of nature of test, frequency of testing and risk factor availability/presence. The main challenge in applying the model to the BVDV CP data was in identifying prior distributions, especially for test characteristics, that corresponded to the latent status of interest, i.e. herds with at least one persistently infected (**PI**) animal. The model is available on Github as an R package (https://github.com/AurMad/STOCfree) and can be used to carry out output-based evaluation of disease CPs.

## Introduction

For many infectious diseases of farm animals, there are control programmes (**CP**s) that rely on the application of diagnostic testing at regular time intervals for the identification of infected animals or herds. In cattle, such diseases notably include infection by the bovine viral diarrhoea virus (**BVDV**) or by *Mycobacterium avium* subspecies *paratuberculosis* (MAP). These CPs are extremely diverse. Their objective can range from decreasing the prevalence of infection to eradication. Participation in the CP can be voluntary or compulsory. The classification of herds regarding infection status can be based on a wide variety of testing strategies in terms of the nature of the tests used (identification of antibodies vs. identification of the agent), the groups of animals tested (e.g. breeding herd vs. young animals), number of animals tested, frequency of testing (once to several times a year, every calf born…). Even within a single CP, surveillance modalities may evolve over time. Such differences in CPs were described by (van Roon, Santman-Berends, et al., 2020) for programmes targeting BVDV infections and by (Whittington et al., 2019) for programmes against MAP.

Differences in surveillance modalities can be problematic when purchasing animals from areas with different CPs because the free status assigned to animals or herds might not be equivalent between CPs. A standardised method for both describing surveillance programmes and estimating confidence of freedom from surveillance data would be useful when trading animals across countries or regions. While inputs can vary between programmes, the output needs to be comparable across programmes. This is called output-based surveillance (Cameron, 2012). Probabilities measure both the chance of an event and the uncertainty around its presence/occurrence. If well designed, a methodology to estimate the probability of freedom from infection would meet the requirements of both providing a confidence of freedom from infection as well as of being comparable whatever the context.

Currently, a common quantitative method used to substantiate freedom from infection to trading partners is the scenario tree method (Martin et al., 2007). The method is applied to situations where there is a surveillance programme in place, with no animals or herds confirmed positive on testing. What is estimated with the scenario tree method is the probability that the infection would be detected in the population if it were present at a chosen *design prevalence*. The output from this approach is the probability that infection prevalence is below the design prevalence given the negative test results (Cameron, 2012). Therefore, this method is well suited for situations where populations are free from infection and those who want to quantify this probability of freedom from infection, e.g. for the benefit of trading partners (Norström et al., 2014).

In a context where disease is controlled but still present, it would only be safe to trade with herds that have an estimated probability of freedom from infection that is deemed sufficiently high or, equivalently, a probability of infection that is deemed sufficiently low. Identifying these herds involves estimating a probability of infection for each herd in the CP and then defining a decision rule to categorise herds as uninfected or infected based on these estimated probabilities.

In this paper, we propose a method to estimate herd level probabilities of infection from heterogeneous longitudinal data generated by CPs. The method predicts herd-month level probabilities of being latent status positive from longitudinal data collected in CPs. The input data are test results, and associated risk factors when available. Our main objective is to describe this modelling framework by showing how surveillance data are related to the *probabilities of infection* (strictly speaking, *probabilities of being latent status positive*) and by providing details regarding the statistical assumptions that are made. A secondary objective is to compare two implementations of this modelling framework, one in JAGS (Plummer, 2003) and one in Stan (Stan Development Team, 2021), for the estimation of these probabilities of being latent status positive. The comparison is performed using surveillance data collected as part of a CP against BVDV infection in Loire-Atlantique, France. The challenges of defining prior distributions and the implications of using different prior distributions are discussed. The functions to perform the analyses described in this paper are gathered in an R package which is available from GitHub (https://github.com/AurMad/STOCfree).

## Materials and methods

### Description of the model

#### Conceptual representation of surveillance programmes

Surveillance programmes against infectious diseases can be seen as imperfect repeated measures of a true status regarding infection. In veterinary epidemiology, the issue of imperfect testing has traditionally been addressed using latent class models. With this family of methods, the true status regarding infection is modelled as an unobserved quantity which is linked to test results through test sensitivity and specificity. Most of the literature on the subject focuses on estimating both test characteristics and infection prevalence (Collins and Huynh, 2014). For the estimations to work, the same tests should be used in different populations (Hui and Walter, 1980), the test characteristics should be the same among populations, and test results should be conditionally independent given the infection status (Johnson et al., 2009; Toft et al., 2005); although some of these assumptions can be relaxed in a Bayesian framework. Latent class models can also be used to estimate associations between infection, defined as the latent class, and risk factors when the test used is imperfect (Fernandes et al., 2019). In the study by (Fernandes et al., 2019), the latent class was defined using a single test, through the prior distributions put on sensitivity and specificity. When using latent class models with longitudinal data, the dependence between successive test results in the same herds must be accounted for. In the context of estimating test characteristics and infection prevalence from 2 tests in a single population from longitudinal data, (Nusinovici et al., 2015) proposed a Bayesian latent class model which incorporated 2 parameters for new infection and infection elimination. The model we describe below combines these different aspects of latent class modelling into a single model.

We propose using a class of models called Hidden Markov Models (HMM, see (Zucchini et al., 2017)). Using surveillance programmes for infectious diseases as an example, the principles of HMMs can be described as follows: the latent status (*class*) of interest is a herd status regarding infection. This status is evaluated at regular time intervals: HMMs are discrete time models. The status at a given time only depends on the status at the previous time (Markovian property). The status of interest is not directly observed, however, there exists some quantity (such as test results) whose distribution depends on the unobserved status. HMMs have been used for decades in speech recognition (Rabiner, 1989) and other areas. They have also been used for epidemiological surveillance (Le Strat and Carrat, 1999; Touloupou et al., 2020), although not with longitudinal data from multiple epidemiological units such as herds. The model we developed is therefore a latent class model that takes into account the time dynamics in the latent status. The probability of new infection between consecutive time steps is modelled as a function of risk factors.

Figure 1 shows how surveillance programmes are represented in the model as a succession of discrete time steps. The focus of this model is a latent status evaluated at the herd-month level. This latent status is not directly observed but inferred from its causes and consequences incorporated as data. The consequences are the test results. Test results do not have to be available at every time step for the model to work, although the estimation will be more accurate with a large number of test results. The causes of infection are risk factors of infection. The model estimates this latent status monthly, and predicts it for the last month of data. These herd-month latent statuses will be estimated/predicted from test results and risk factors recorded in each herd.

**Figure 1.**
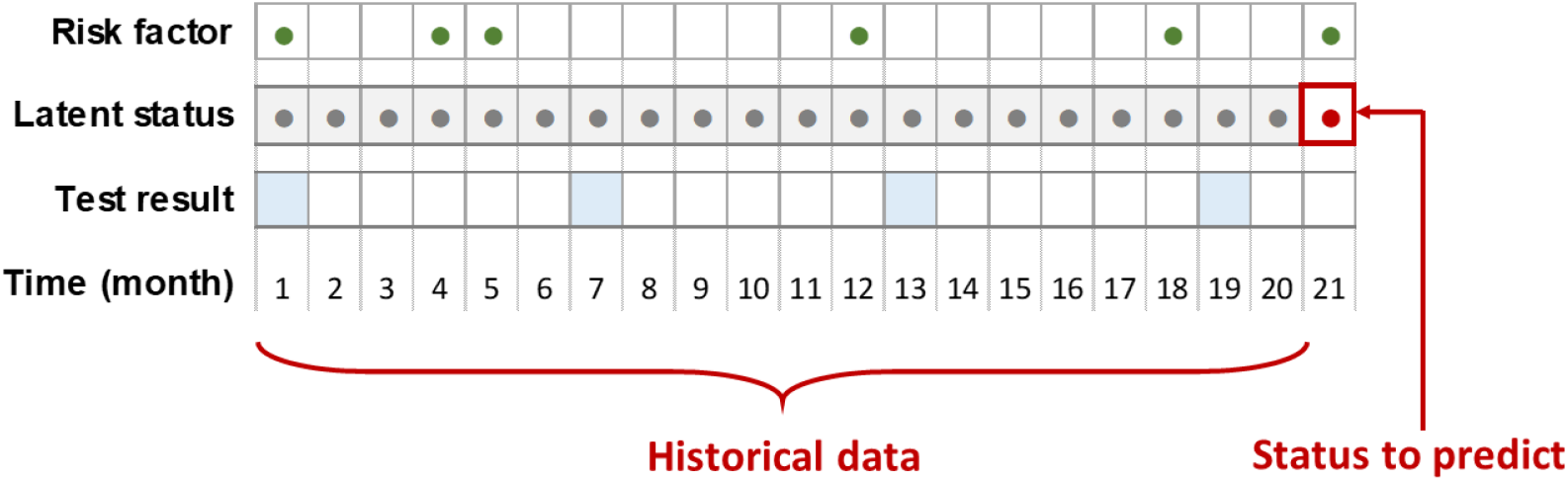
Conceptual representation of the implementation of a surveillance programme within a herd. The focus of the model is the latent status regarding infection, which is modelled at the herd-month level. This status partly depends on risk factors and determines test results. In this diagram, risk factors are represented as green dots when present and available test results as blue shaded squares. The model predicts a probability of infection for the most recent month in the surveillance programme using all the data collected for the estimation of model parameters.

#### Modelling framework, inputs and outputs

The model is designed to use longitudinal data collected as part of surveillance programmes against infectious diseases. In such programmes, each herd level status is re-evaluated when new data (most commonly test results, but may also be data related to risk factors) are available. The model mimics this situation by predicting the probability of a positive status for all herds in the CP on the last month of available data. Data from all participating herds up to the month of prediction are used as historical data for parameter estimation (Figure 1).

The estimation and prediction are performed within a Bayesian framework using Markov Chain Monte Carlo (MCMC). The model encodes the relationships between all the variables of interest in a single model. Each variable is modelled as drawn from a statistical distribution. The estimation requires prior distributions for all the parameters in the model. These priors are a way to incorporate either existing knowledge or hypotheses in the estimation. For example, we may know that the prevalence of herds infected with BVDV in our CP is probably lower than 20%, certainly lower than 30% and greater than 5%. There are different ways of specifying such constraints using statistical distributions. We will briefly describe two that are used in different places in our modelling framework. The first one consists in using a Beta distribution. The Beta distribution is bounded between 0 and 1, with 2 parameters *α* and *β* determining its shape. With the constraints specified above, we could use as a prior distribution *Beta*(*α* = 15, *β* = 100)^1^. The second one consists in using a normal distribution on the logit scale. The principle of the logit transformation is to map probabilities that are bounded between 0 and 1 onto an interval that extends from −∞ to +∞. Quantities defined on the logit scale, can be mapped back onto the probability scale using the inverse logit transformation ^2^. This is extremely convenient because it allows the use of normal distributions on the logit scale, whose mean and standard deviation have an intuitive meaning. With the constraints specified above, we could use as a prior distribution a *Normal*(*μ* = −2, *σ*^2^ = 0.09) ^3^. If we do not know anything about this infection prevalence (which is rare), we could use a *Beta*(*α* = 1, *β* = 1) prior, which is uniform between 0 and 1; or a *Normal*(*μ* = 0, *σ*^2^ = 10) on the logit scale. From the model specification, the prior distributions and the observed data, the MCMC algorithm draws samples from the posterior distributions of all the variables in the model. These posterior distributions are the probability distributions for the model parameters given the data and the prior distributions. MCMC methods are stochastic and iterative. Each iteration is a set of samples from the joint posterior distributions of all variables in the model. The algorithm is designed to reach the target joint posterior distribution, but at any moment, there is no guarantee that it has done. To overcome this difficulty, several independent instances of the algorithm (i.e. several chains) are run in parallel. For a variable, if all the MCMC draws from the different chains are drawn from the same distribution, it can be concluded that the algorithm has reached the posterior distribution. In this case, it is said that the model has converged.

The focus of our model is the monthly latent status of each herd. This latent status depends on the data on occurrence of risk factors and it affects test results. The data used by the model are the test results and risk factors. At each iteration of the MCMC algorithm, given the data and priors, a herd status (0 or 1) and the coefficients for the associations between risk factors, latent status and test results are drawn from their posterior distribution.

In the next 3 sections, the parameters for which prior distributions are required, i.e. test characteristics, status dynamics and risk factor parameters, are described. The outputs of Bayesian models are posterior distributions for all model parameters. Specifically, in our model, the quantities of interest are the herd level probabilities of being latent status positive on the last test month in the dataset as well as test sensitivity, test specificity, infection dynamic parameters and parameters for the strengths of association between risk factors and the probability of new infection. This is described in the corresponding sections.

#### Latent status dynamics

Between test events, uninfected herds can become infected and infected herds can clear the infection. The model represents the probability of having a positive status at each time step as a function of the status at the previous time step (Figure 2). For the first time step when herd status is assigned, there is no previous status against which to evaluate change. From the second time step when herd status is assigned, and onwards, herds that were status negative on the previous time step have a certain probability of becoming status positive and herds that were status positive have a certain probability of remaining status positive.

**Figure 2.**
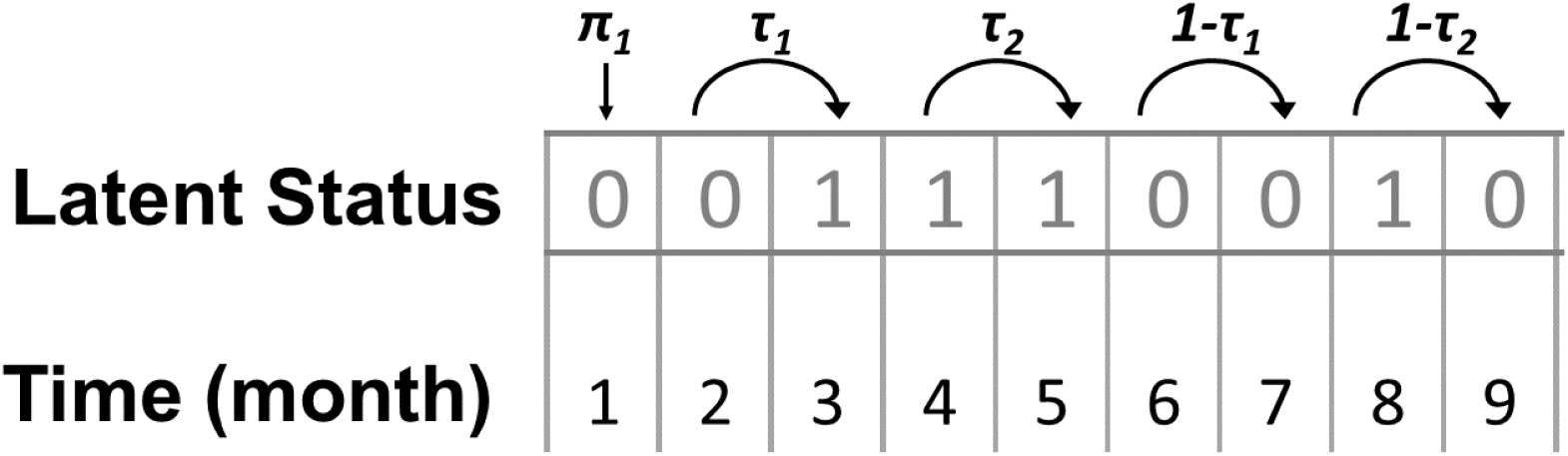
Modelling of infection dynamics. The diagram shows hypothetical latent statuses (0 for negative; 1 for positive) as a function of time in month, with examples of all possible transitions. *π*_1_ = *p*(*S*_1_ = 1) is the probability of being status positive at the first point in time, *τ*_1_ = *p*(*S*_*t*_ = 1|*S*_*t−*1_ = 0) is the probability of becoming status positive and *τ*_2_ = *p*(*S*_*t*_ = 1|*S*_*t−*1_ = 1) is the probability of remaining status positive.

These assumptions can be summarised with the following set of equations^4^. The status on the first time step (*S*_1_) is a Bernoulli event with a normal prior on the logit scale for its probability of occurrence:

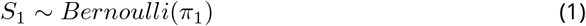

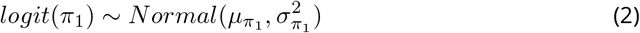

From the second time step when herd status is assigned, and onwards, a positive status is also a Bernoulli event (*S*_*t*_) with a probability of occurrence that depends on the status at the previous time step as well as on the probability of becoming status positive and the probability of remaining status positive. In this case, the probability of becoming status positive is *τ*_1_ = *p*(*S*_*t*_ = 1|*S*_*t−*1_ = 0) and the probability of remaining positive is *τ*_2_ = *p*(*S*_*t*_ = 1|*S*_*t−*1_ = 1).

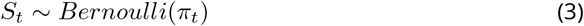

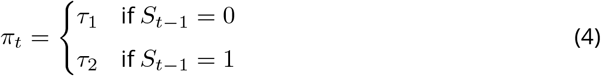

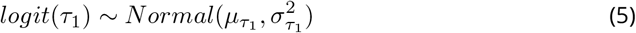

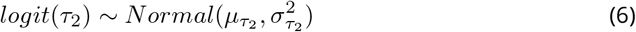

Therefore, the status dynamics can be completely described by *π*_1_, *τ*_1_ and *τ*_2_.

#### Incorporation of information on risk factors for new infection

The probability of new infection is not the same across herds. For example, herds that introduce a lot of animals or are in areas where infection prevalence is high could be at increased risk of new infection (Qi et al., 2019). Furthermore, the association between a given risk factor and the probability of new infection could be CP dependent. For example, the probability of introducing infection through animal introductions will depend on the infection prevalence in the population from which animals are introduced. As a consequence, estimates for these associations (as presented in the literature) could provide an indication about their order of magnitude, but their precision may be limited. On the other hand, the CPs which are of interest in this work usually generate large amounts of testing data which could be used to estimate the strengths of association between risk factors and new infections within a given CP. The variables that are associated with the probability of new infection could increase the sensitivity and timeliness of detection.

When risk factors for new infection are available, the model incorporates this information by modelling *τ*_1_ as a function of these risk factors through logistic regression, instead of the prior distribution for *τ*_1_.

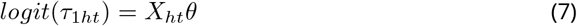

where *X*_*ht*_ is a matrix of predictors for herd *h* at time *t* and *θ* is a vector of coefficients. Normal priors are used for the coefficients of the logistic regression.

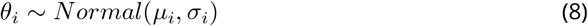

#### Test characteristics

The model allows the inclusion of several test types but for the sake of clarity, we show the model principles for only one test type. These principles can be extended to several tests by specifying prior distributions for all tests.

Tests are modelled as imperfect measures of the latent status (Figure 3). Test sensitivity is the probability of a positive test result given a positive latent status (*Se* = *p*(*T* = 1|*S* = 1), refers to true positives) and test specificity is the probability of a negative test result given a negative latent status (*Sp* = *p*(*T* = 0|*S* = 0), refers to true negatives).

**Figure 3.**
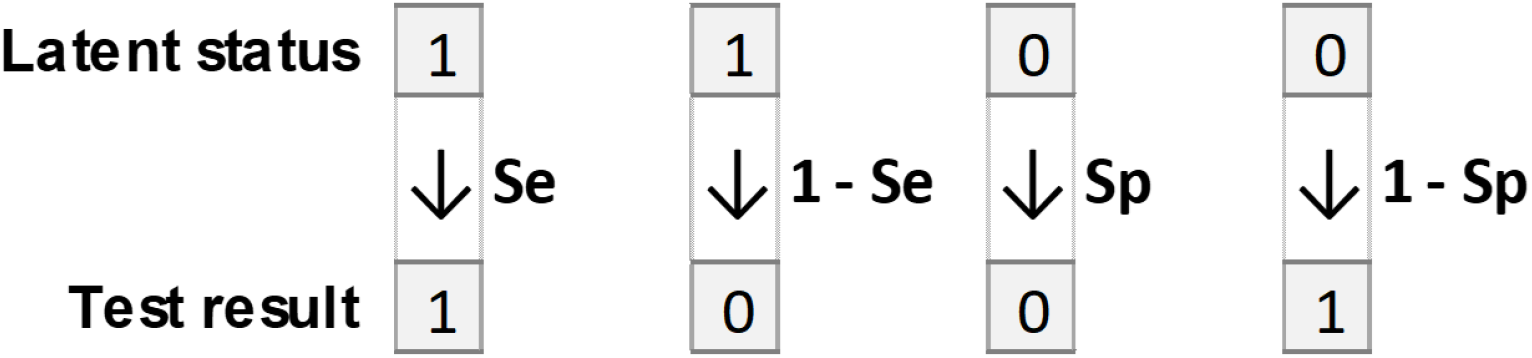
Relation of the model latent status to test result. Sensitivity is the probability of a positive test result in a status positive herd. Specificity is the probability of a negative test result in a status negative herd.

Test result at time *t* is modelled as a Bernoulli event with probability *p*(*T*_*t*_) of being positive.

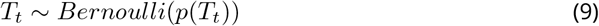

The relation between the probability of testing positive, the probability of a positive status, test sensitivity and test specificity is the following:

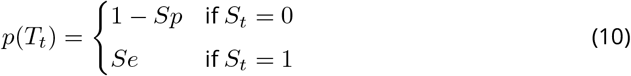

Information or hypotheses regarding test characteristics are incorporated in the model as priors modelled by Beta distributions:

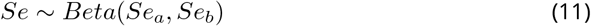

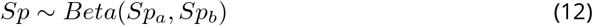

#### Prediction of a probability of infection in JAGS

In JAGS, a specific step was needed in order to predict the final probability of being status positive given historical data and a test result on the month of prediction, when such a test result was available. In Stan, this step was not necessary because the forward algorithm directly predicted the probability of being status positive in the last month. In explaining how predictions are performed in JAGS, we use the following notation: 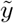 is the predicted value for 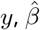 is the estimated value for *β*. The equation 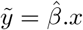 means that the predicted value for *y* is equal to *x* (data) times the estimated value for *β*.

The model predicts herd-level probabilities of being latent status positive on the last month in the data mimicking regular re-evaluation as new data come in. If there is no test result available on this month, the predicted probability of being status positive 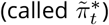 is the predicted status on the previous month times 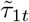 if the herd was predicted status negative or times 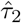 if the herd was predicted status positive (Table 1)^5^. This can be written as:

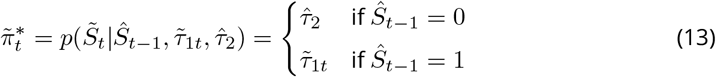

where:

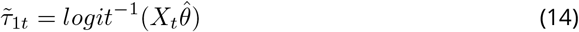

**Table 1.** Probability of test result by herd status. Cells on the first row are test positive herds with true positives on the left-hand side and false positives on the right-hand side. Cells on the second row are test negative herds with false negatives on the left-hand side and true negatives on the right-hand side.

|  |  | Herd status<sub>t</sub> |  |
| --- | --- | --- | --- |
|  |  | + | - |
| Test<sub>t</sub> | + | $Se.\pi_t$ | $(1 - Sp)(1 - \pi_t)$ |
| | - | $(1 - Se).\pi_t$ | $Sp.(1 - \pi_t)$ |

If a test result was available, the prediction must combine information from the test as well as previous information. The way to estimate this predicted probability from 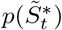 and test results can be derived from Table 1. The predicted probability of being status positive can be computed as:

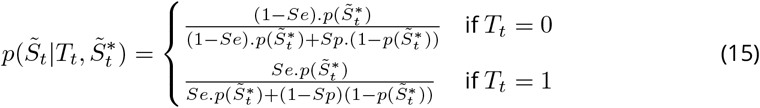

where *T*_*t*_ = 1 when the test at time *t* is positive, *T*_*t*_ = 0 when it is negative

#### Model implementations

The pre-processing of the data and the analysis of the results of the Bayesian models were done in R (R Core Team, 2020). The HMM was implemented in both JAGS and Stan.

The model was initially implemented in JAGS, which performs Bayesian inference using Gibbs sampling (Plummer, 2003). The model equations were directly translated into JAGS code. The `runjags` R package (Denwood, 2016) was used to interface R and JAGS.

The model was then implemented in Stan (Stan Development Team, 2021). Stan is a newer and more efficient way of performing Bayesian inference using Hamiltonian Monte Carlo. However, Stan does not allow latent discrete parameters to be modelled directly. Therefore, for the Stan implementation of our model, the forward algorithm (Baum and Eagon, 1967) was adapted from (Damiano et al., 2018). The `cmdstanr` R package (Gabry and Cešnovar, 2020) was used to interface R and Stan.

### Application of the model to a control programme for BVDV infection in cattle

#### Data

The model was evaluated on data collected for the surveillance of BVDV infection in dairy cattle in Loire-Atlantique, France. Under the programme, each herd was tested twice a year with a bulk tank milk (**BTM**) antibody ELISA test. For each campaign of testing, tests were performed for all herds over a few weeks. Data on the number of cattle introduced into each herd with the associated date of introduction were also available. For the model evaluation, test data of 1687 herds from the beginning of 2014 to the end of 2016 were used. Risk factor data collected between 2010 and 2016 were available to model (possibly lagged) associations between risk factors and the latent status.

#### Test results

Test results were reported as optical density ratios (ODR). These ODR values were discretised in order to convert them into either seropositive (antibodies detected) or seronegative (no antibodies detected) outcomes. The choice of the threshold to apply for the discretisation as well as the sensitivity and specificity of this threshold for the detection of seropositivity were based on the ODR distributions from test data collected outside of the study period. The overall ODR distribution was modelled as a mixture of underlying ODR distributions for seropositives and seronegtaives. The details of the method used are provided as supplementary material.

#### Selection of risk factors

A difficulty in the evaluation of putative risk factors was that Bayesian models usually take time to run, especially with large datasets as used here. It was therefore not possible to perform this selection with our Bayesian model. To circumvent this problem, logistic models as implemented in the R glm function (R Core Team, 2020) were used^6^. The outcome of these models was seroconversion defined as a binary event, and covariates of interest were risk factors for becoming status positive as defined through the *τ*_1_ variable. All herds with 2 consecutive test results whose first result was negative (ODR below the chosen threshold) were capable of seroconverting. Of these herds, the ones that had a positive result (ODR above the chosen threshold) on the second test were considered as having seroconverted. The time of event (seroconversion or not) was considered the mid-point between the 2 tests.

Two types of risk factors of new infection were evaluated: infection through cattle introductions and infection through neighbourhood contacts (Qi et al., 2019). Cattle introduction variables were constructed from the number of animals introduced into a herd on a given date. In addition to the raw number of animals introduced, the natural logarithm of the number of animals (+1 because ln(0) is not defined) was also evaluated. This was to allow a decreasing effect of each animal as the number of animals introduced increased. Regarding the neighbourhood risk, the test result data were used. For each testing campaign, the municipality-level prevalence of test positives (excluding the herd of interest) was calculated, and is subsequently termed ‘local prevalence’. It was anticipated that when local seroprevalence would increase, the probability of new infection in the herd of interest would increase as well.

For all candidate variables, a potential problem was delayed detection, which relates to the fact that a risk factor recorded at one point in time may be detected through testing much later, even if the test is sensitive. For example, if a trojan cow (a non-PI female carrying a PI calf) is introduced into a herd, the lactating herd will only seroconvert when the PI calf is born and has had contact with the lactating herd. Therefore, for each candidate variable, the data were aggregated between the beginning of an interval (labelled lag1, in months from the outcome measurement) and the end of this interval (labelled lag2, in months from the outcome measurement). Models with all possible combinations of time aggregation between lag1 and lag2 were run, with lag1 set to 0 and lag2 set to 24 months. The best variables and time aggregation interval were selected based on low AIC value, biological plausibility and suitability for the Bayesian model.

#### Bayesian models

Four different Bayesian models were considered. For all models, historical data were used for parameter estimation and the probability of infection on the last month in the dataset was predicted.

##### Model 1 - Perfect test, no risk factors

in order to evaluate the monthly dynamics of seropositivity and seronegativity, the Bayesian model was run without any risk factors and assuming that both test sensitivity and test specificity were close to 1. The prior distributions for sensitivity and specificity were *Se* ∼ *Beta*(10000, 1) (percentiles: 5 = 1, 50 = 1, 95 = 1) and *Sp* ∼ *Beta*(10000, 1). Regarding infection dynamics, prior distributions were specified for the prevalence of status positives (also test positives in this scenario) on the first testing 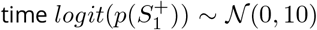. (on the probability scale - percentiles: 5 = 0, 50 = 0.5, 95 = 1), the probability of becoming status positive *logit*(*τ*_1_) ∼ 𝒩(−3, 1) (percentiles: 5 = 0.01, 50 = 0.047, 95 = 0.205), and the probability of remaining status positive *logit*(*τ*_2_) ∼ 𝒩(2.2, 0.05) (percentiles: 5 = 0.893, 50 = 0.9, 95 = 0.907). The same prior distribution for *τ*_2_ was used in all models. The motivation for this choice was the fact that tests were performed every 6 months in all herds. The consequences of choosing this prior was that infected herds had a small probability of changing status between consecutive months (median probability = 0.1), but after 6 months, the probability of still being positive was 0.9^6^ = 0.53, at which time the status was updated with a new test result.

##### Model 2 - Imperfect test, no risk factors

the objective of this model was to incorporate the uncertainty associated with test results in both parameter estimation and in the prediction of the probabilities of infection. The priors for test sensitivity and specificity were selected based on the ODR distributions for seronegatives and seropositives identified by the mixture model. The following prior distributions were used: *Se* ∼ *Beta*(10, 1) (percentiles: 5 = 0.741, 50 = 0.933, 95 = 0.995) and *Sp* ∼ *Beta*(10, 1). For the status dynamics parameters, the same prior distributions as in Model 1 were used.

##### Model 3 - Perfect test, risk factors

in order to quantify the association between risk factors and the probability of becoming status positive if the test were close to perfect, the Bayesian model was run with the risk factors identified as associated with seroconversion on the previous step, and using the same priors for sensitivity, specificity and *τ*_2_ as in Model 1. The priors for risk factors were specified as normal distributions on the logit scale. The prior for the intercept was *θ*_1_ ∼ 𝒩(−3, 1) (on the probability scale - percentiles: 5 = 0.01, 50 = 0.047, 95 = 0.205). This represented the prior probability of a new infection in a herd purchasing no animal and with a local seroprevalence of 0. The priors for the other model coefficients were centred on 0 with a standard deviation of 2. On the logit scale, values of -4 (2 standard deviations in this case) correspond to probabilities close to 0 (logit(−4) = (0.018) and values of 4 to probabilities that are close to 1 (logit(4) = (0.982).

##### Model 4 - Imperfect test, risk factors

in order to quantify the association between risk factors and the probability of becoming status positive while incorporating test imperfection, the Bayesian model was run with the risk factors identified as associated with seroconversion using the same priors as in Model 1 for tests characteristics and the same priors as in Model 3 for infection dynamics and risk factors.

Each model was run in both Stan and JAGS. For each model, 4 chains were run in parallel. For the Stan implementation, the first 1 000 iterations were discarded (warmup). The model was run for 500 more iterations with every iteration stored for analysis. This yielded 2 000 draws from the posterior distribution of each parameter. For the JAGS implementation, the first 15 000 MCMC iterations were discarded (burn-in). The model was run for 10 000 more iterations of which 1 in 20 was stored for analysis. This yielded 2 000 draws from the posterior distribution of each parameter. For all models, convergence was assessed visually using traceplots. Each distribution was summarised with its median and 95% credibility interval.

## Results

### Test results

Between the beginning of 2014 and the end of 2016, there were 9725 available test results, reported as ODRs, from 1687 herds. Most herds were tested in February and September (See Figure 4). The cut-off of 35 used in the CP seemed to discriminate well between the distributions associated with seronegative and seropositive herds respectively, and was therefore retained in the remainder of the analysis. Using this threshold, there were 44.1% of seropositive tests between 2014 and 2016. The associated estimated test sensitivity and specificity were 0.978 and 0.949 respectively. However, in the Bayesian models 2 and 4, because there was considerable uncertainty regarding the assumptions made, sensitivity and specificity were modelled using *Beta*(10, 1) prior distributions (percentiles: 5 = 0.741, 50 = 0.933, 95 = 0.995).

**Figure 4.**
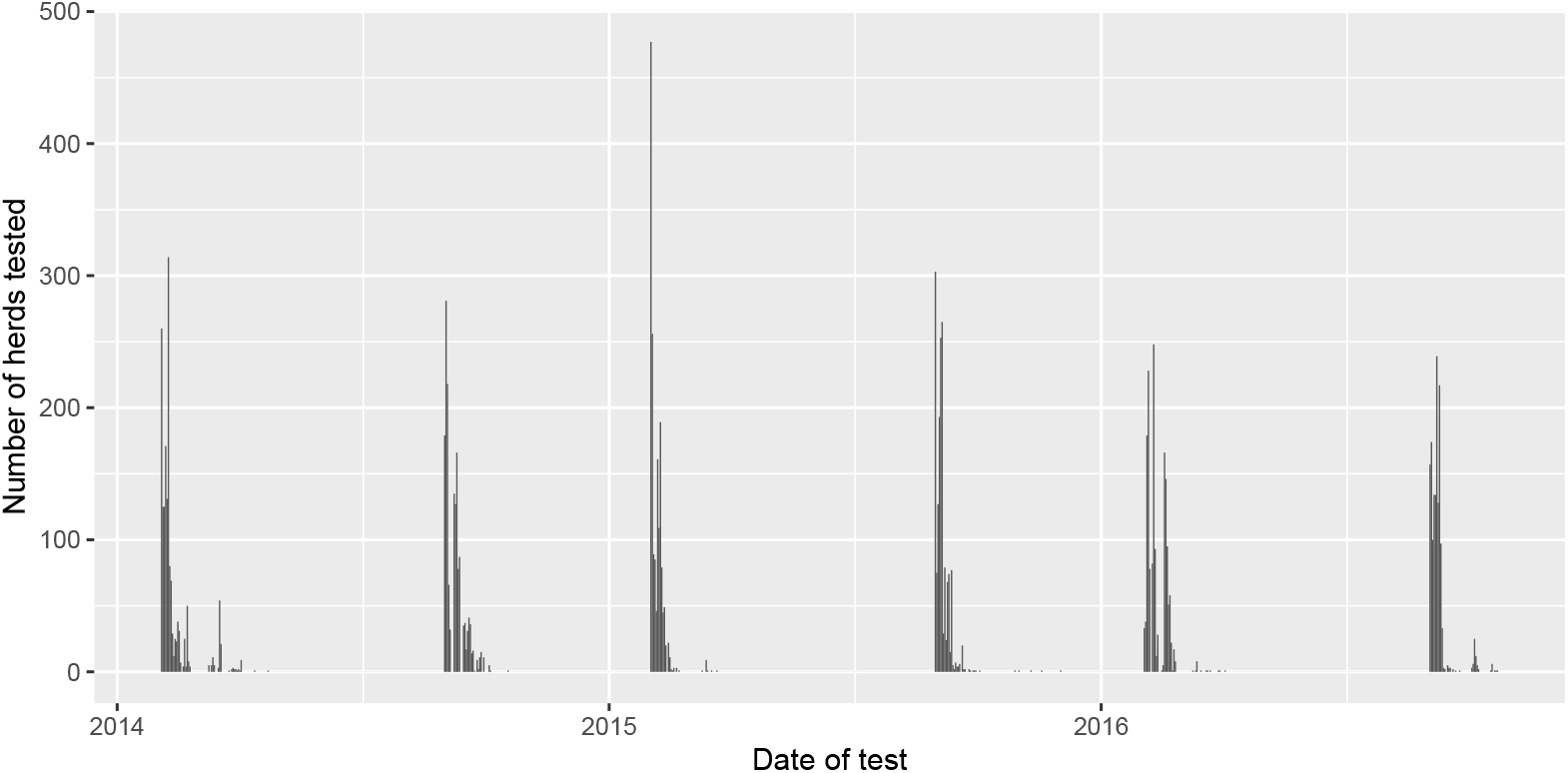
Distribution of the test dates between 2014 and 2017 in 1687 herds from Loire-Atlantique, France.

### Selection of risk factors

Risk factors related to animal introductions and seroprevalence were evaluated with logistic models. The model outcome was a seroconversion event. A first step of the analysis was, for each variable, to identify the time interval that was the most predictive of an observed seroconversion. Figure 5 presents the AIC values associated with each possible interval for the variables ln(Number of animals introduced + 1) and local seroprevalence.

**Figure 5.**
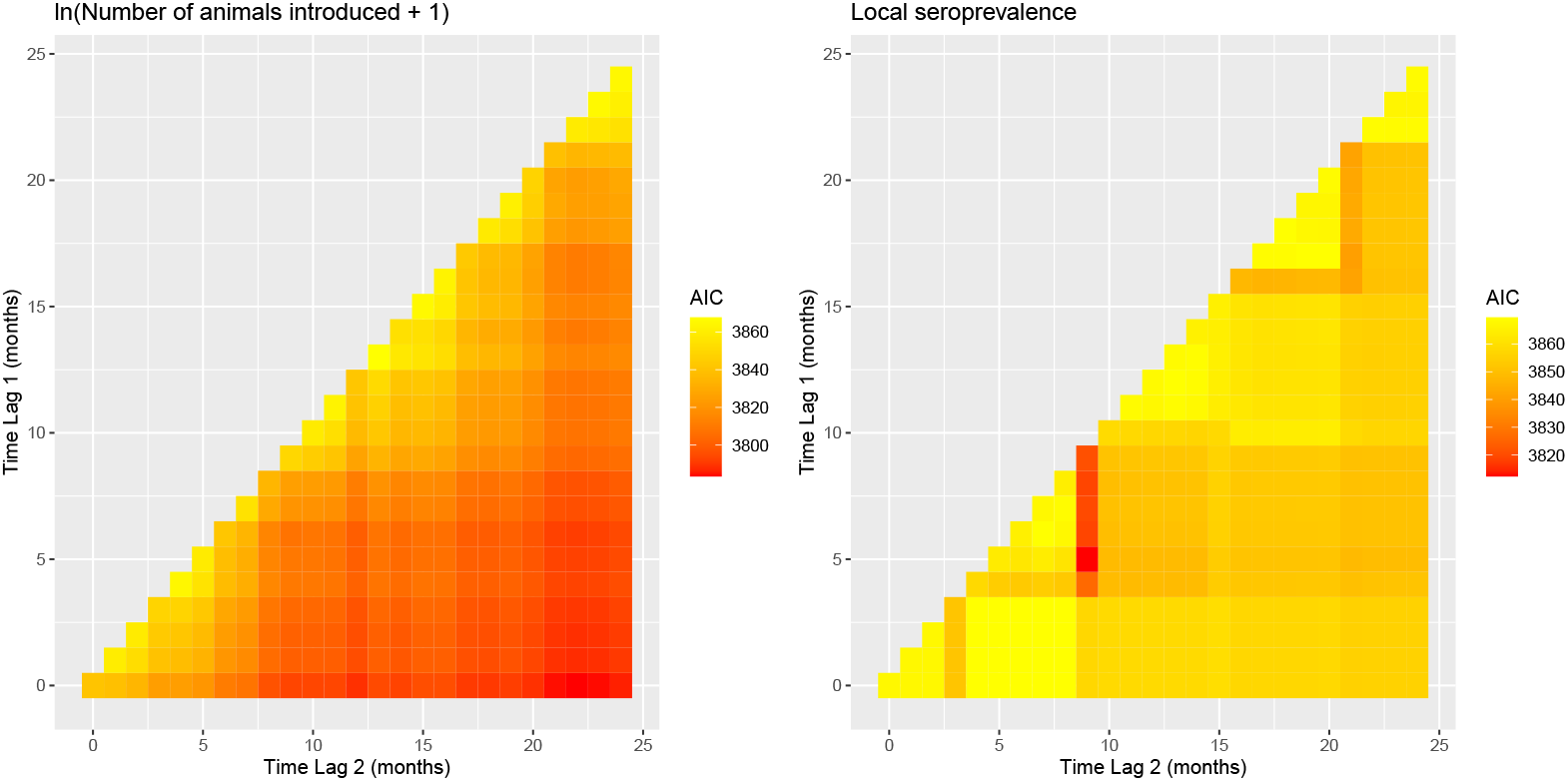
AIC values associated with logistic models of the association between 2 variables and the probability of seroconversion between 2 tests. Each coloured row represents the end of the interval that is closest to the current month and each column the end of the interval that is the furthest in the past. For example, the line at the bottom represents intervals that end on the current month: the first column is for the interval that started and ended on the current month (length of one month) and the last column is for the interval between 24 months ago and the current month. The variable evaluated on the left-hand side panel is the sum of the ln(number of animals introduced + 1) between lag1 and lag2. The variable evaluated on the right-hand side panel is the max of the local seroprevalence between lag1 and lag2.

For the animal introduction variables, for the same time interval, the AICs of the models of the untransformed number of animals were higher than the ones for the log transformed values (not shown). It can also be noted that considering longer intervals (further away from the diagonal) was usually better than considering short intervals (close to the diagonal). It may be that some herds never buy any animal while, on average, herds that buy once have already done it in the past. In this case, it is possible that the infection was introduced several times, while it is not possible to know which animal introduction was associated with herd seroconversion. This could explain the apparent cumulative effect of the number of introductions. The cells that are close to the diagonal are associated with short intervals. Considering one month intervals, the probability of infection was highest for introductions made 8 months from the month of seroconversion.

Local seroprevalence was evaluated from data collected in 2 different testing campaigns per year, as shown in Figure 4. For this reason, in the investigation of lagged relationships between local seroprevalence and the probability of seroconversion, the maximum local seroprevalence was computed, and not the sum as for the number of animals introduced. The strength of association between local seroprevalence and herd seroconversion was greatest for local seroprevalence 9 months prior to herd seroconversion.

A final multivariable logistic model with an animal introduction variable and a local sero-prevalence variable was constructed. In the choice of the time intervals to include in this model, the following elements were considered. First, the Bayesian model runs with a monthly time step. Aggregating data over several months would result in including the same variable several times. Secondly, historical data may sometimes be limited. Having the smallest possible value for the end of the interval could be preferable. For this reason, the variables considered for the final model were the natural logarithm of the number of animals introduced 8 months prior to the month of seroconversion as well as the local seroprevalence 9 months prior to the month of seroconversion. The results of this model are presented in Table 2. All variables were highly significant. The model intercept was the probability of seroconversion in a herd introducing no animals and with local seroprevalence of 0 in each of the time intervals considered. The probability of seroconversion between 2 tests corresponding to this scenario was of 0.124. Buying 1, 10 or 100 animals increased this estimated probability to 0.171, 0.866 and 1 respectively. Buying no animals and observing a seroprevalence of 0.2 (proportion of seropositives in the dataset) was associated with a probability of seroconversion of 0.261.

**Table 2.** Results of the final logistic model of the probability of seroconversion between consecutive tests. The risk factors retained in the model were the logarithm of the number of animals introduced in the herd 8 months before seroconversion and the local seroprevalence 9 months before seroconversion.

|  | lag1 | lag2 | Estimate | p-value |
| --- | --- | --- | --- | --- |
| Intercept | - | - | -1.96 | < 0.001 |
| ln(Number animals introduced +1) | 8 | 8 | 0.38 | < 0.001 |
| local seroprevalence | 9 | 9 | 4.59 | < 0.001 |

### Bayesian models

Running the different models for the 1687 herds with 3 years of data on the first author’s laptop (CPU: Intel Core i5-8350U, RAM: 16 Go, Windows 10) took significantly more time in JAGS (3 to 4.5 hours) than in Stan (around 1 hour). In models 3 and 4, the candidate covariates were the natural logarithm of the number of animals introduced 8 months before status evaluation/prediction as well as the local seroprevalence 9 months before. The 95% credibility interval for the estimated coefficient associated with local seroprevalence included 0. This variable was therefore removed from the models and only cattle introductions were considered.

#### Model parameters

For Models 1 and 3, in which the test was assumed to be perfect, the 4 chains of each model converged and mixed well regardless of the programme used for Bayesian inference. For Models 2 and 4, in which wider distributions were assumed for test characteristics, the chains converged and mixed well for the Stan version, but mixing was poor for the JAGS version. As an illustration, Figure 6 represents the traceplots for test sensitivity in Models 1 and 2 with both the Stan and JAGS version of the models. In the JAGS version of Model 2, autocorrelation is visible in the traceplot for sensitivity, despite the fact that only one iteration in 20 (thinning of 20) was kept for analysis. Figure 7 and Table 3 show the distributions of model parameters for the 4 models. Although the JAGS model tends not to converge as well, the parameter estimates are similar between the Stan and JAGS versions of the models.

**Table 3.** Median (2.5%, 97.5%) of the parameter posterior distributions used in the 4 Bayesian models evaluated. Model 1: Perfect routine test; Model 2: Perfect routine test and risk factors; Model 3: Imperfect routine test and risk factors; Model 4: Imperfect routine test, confirmatory testing and risk factors.

| Model | Inference | Se | Sp | $\tau_1 / \logit^{-1}\theta_1$ | $\theta_2$ | $\tau_2$ |
| --- | --- | --- | --- | --- | --- | --- |
| Model 1 | Stan | 1 (1-1) | 1 (1-1) | 0.032 (0.03-0.034) | - | 0.948 (0.945-0.951) |
|  | JAGS | 1 (1-1) | 1 (1-1) | 0.032 (0.03-0.034) | - | 0.948 (0.945-0.951) |
| Model 2 | Stan | 0.982 (0.968-0.994) | 0.946 (0.934-0.957) | 0.018 (0.016-0.021) | - | 0.958 (0.954-0.96) |
|  | JAGS | 0.967 (0.953-0.982) | 0.954 (0.943-0.965) | 0.018 (0.015-0.02) | - | 0.958 (0.955-0.961) |
| Model 3 | Stan | 1 (1-1) | 1 (1-1) | 0.028 (0.026-0.031) | 0.615 (0.508-0.716) | 0.948 (0.945-0.95) |
|  | JAGS | 1 (1-1) | 1 (1-1) | 0.028 (0.026-0.031) | 0.613 (0.508-0.721) | 0.947 (0.944-0.95) |
| Model 4 | Stan | 0.979 (0.966-0.989) | 0.948 (0.937-0.959) | 0.015 (0.013-0.018) | 0.725 (0.596-0.842) | 0.957 (0.954-0.96) |
|  | JAGS | 0.969 (0.956-0.982) | 0.955 (0.943-0.965) | 0.015 (0.013-0.018) | 0.731 (0.606-0.856) | 0.957 (0.954-0.96) |

**Figure 6.**
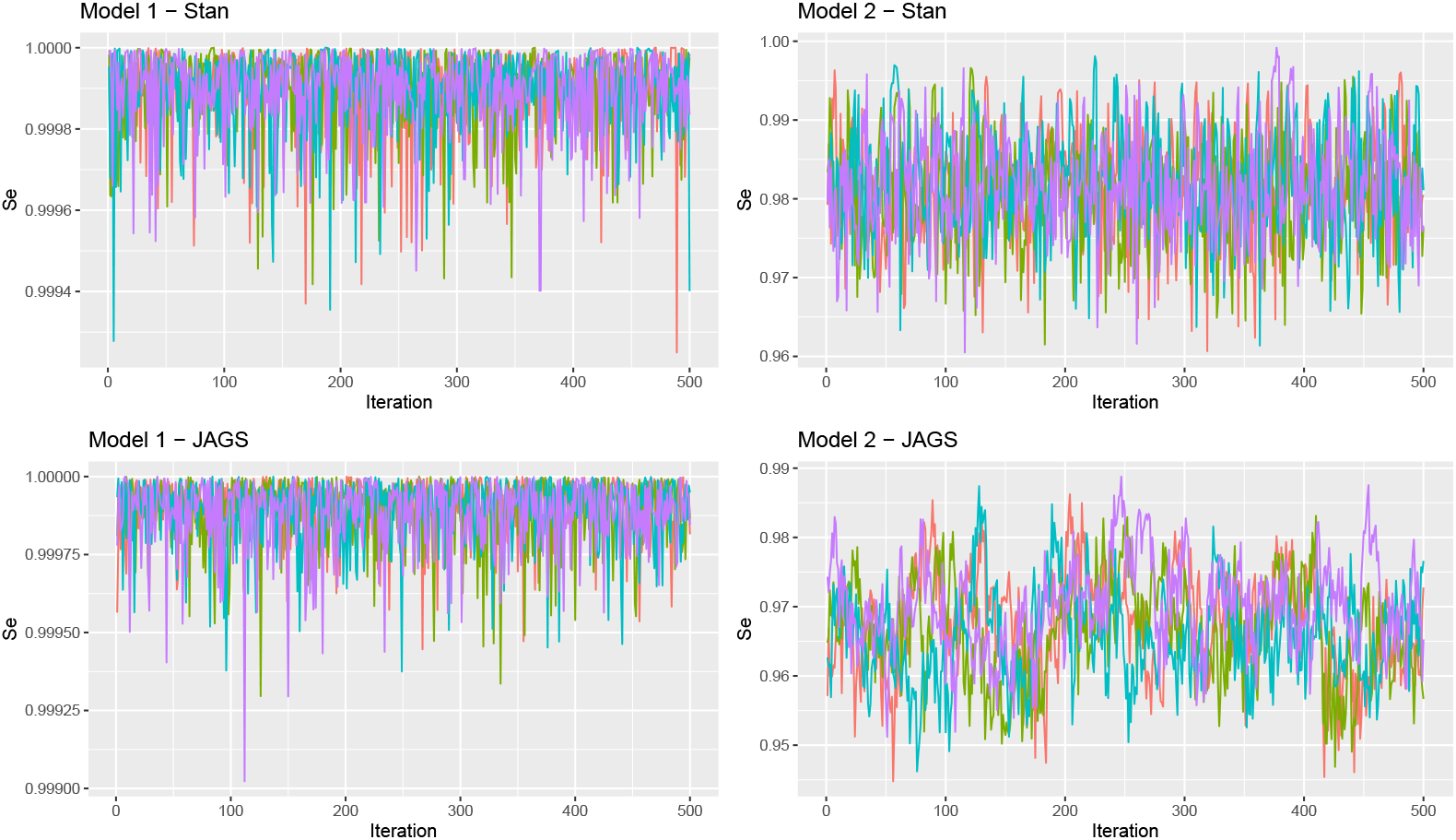
Traceplots for test sensitivity in Models 1 and 2 estimated in Stan and JAGS. Each color represents one of 4 chains run for each model.

**Figure 7.**
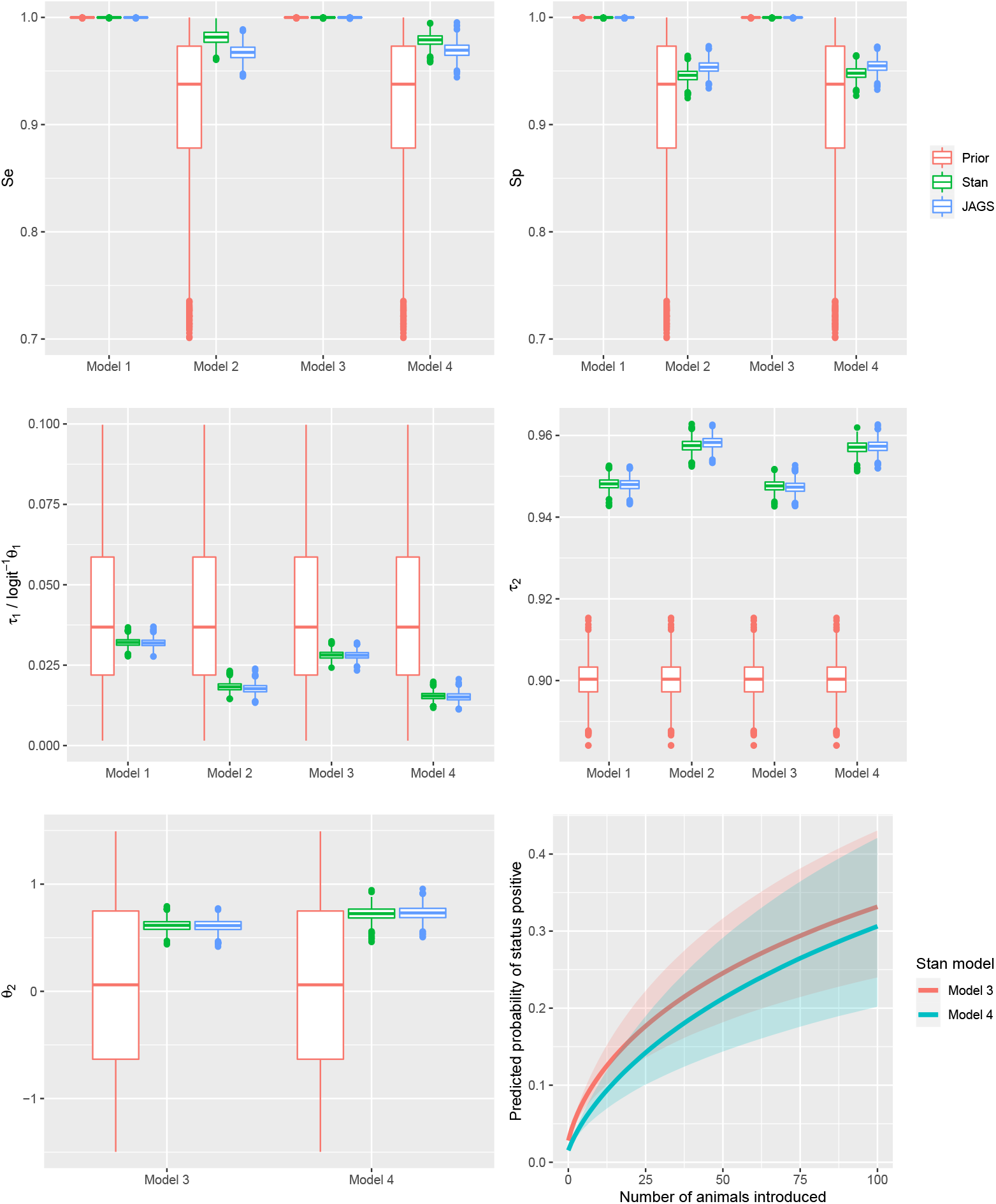
Parameters prior and posterior distributions for the 4 Bayesian models. Model 1: Perfect test, no risk factor; Model 2: Imperfect test, no risk factor; Model 3: Perfect test, risk factor; Model 4: Imperfect test, risk factor. The only risk factor included is the logarithm of the number of animals introduced + 1. In Models 1 and 2, the probability of becoming status positive is modelled with *τ*_1_. In Models 3 and 4, the probability of becoming positive is modelled using logistic regression. From these models, *logit*^−1^ *τ*_1_ is the probability of becoming positive when no animal is introduced (i.e. model intercept). *τ*_2_ models the increase in the probability of becoming positive with the number of animals introduced. The last row of the Figure represents the posterior distribution for *τ*_2_ as well as the corresponding increase in the probability of becoming positive with the number of animals introduced.

In Model 1, the prior distribution put on sensitivity and specificity was very close to 1. With this model, the latent status corresponded to the test result. In effect, it modelled the monthly probability of transition between BTM test negative and BTM test positive. In this case, the median (percentile 2.5 - percentile 97.5) probability of becoming status positive between consecutive months was 0.032 (0.030 - 0.034). This represents a probability of becoming status positive over a 12 month period of 0.323 (0.310 - 0.340). For status positive herds, the monthly probability of remaining positive was of 0.948 (0.945 - 0.951) which represents a probability of still being status positive 12 months later of 0.526 (0.507 - 0.547).

In models 2 and 4, a *Beta*(10, 1) distribution was used as a prior for test sensitivity and specificity. Despite this distribution spanning a relatively large interval (percentiles: 5 = 0.741, 50 = 0.933, 95 = 0.995), all models converged to high values for both sensitivity and specificity. As noted above, convergence was not as good for the JAGS versions of the models, although the JAGS and Stan estimates are close. Interestingly, for model parameters related to status dynamics and risk factors, the Stan and JAGS estimates were almost identical for all models. Adding test imperfection to the models resulted in a decrease in the probability of becoming positive (from 0.032 to 0.018 between models 1 and 2; from 0.028 to 0.015 between models 3 and 4) as well as in an increase in the probability of remaining positive (from 0.948 to 0.958 between models 1 and 2; from 0.948 to 0.957 between models 3 and 4). The most likely reason is that, in some herds, some negative tests arising in a sequence of positive tests were considered as false negatives resulting in longer sequences of positive status and, as a consequence, fewer transitions from negative to positive status.

In models 3 and 4, a risk factor of becoming status positive was incorporated into the estimation. The model intercept (*θ*_1_) was much lower than the estimate from the logistic model estimated in the variable selection step. This was due to the different time steps considered (1 month vs. half a year). On the other hand, the estimate for the association between the natural logarithm of the number of animals introduced and the probability of becoming positive was higher. This association is plotted in the bottom right-hand side panel of Figure 7. The probability of becoming latent status positive between 2 months goes from 0.015 when introducing no animal (*logit*^*−*1^*θ*_1_ in Table 3) to greater than 0.3 for 100 animals introduced. This suggests that including the number of animals introduced into the prediction of herd statuses could increase the sensitivity of detection.

### Predicted probabilities of infection

Figure 8 shows the distributions of herd-level probabilities of infection predicted by the 4 Bayesian models, using Stan and JAGS. These probability distributions are bimodal for all models. The left-hand side corresponds to herds that were predicted status negative on the month before the month of prediction. These are associated to becoming status positive, i.e. *τ*_1_. The right-hand side of the distributions corresponds to herds that were predicted status positive on the month before the month of prediction. These are associated to remaining status positive, i.e. *τ*_2_. Figure 9 shows the distributions of the predicted probability of being status positive for 4 herds. It can be seen that herds that were consistently negative (positive) to the test had extremely low (high) probabilities of being status positive. Accounting for the number of animals introduced increased the probability of infection in the herds that were test negative. An important difference between JAGS and Stan was that in JAGS latent statuses are explicitly represented as a binary variable. As a consequence, herds can *jump* between status positive and status negative on the month before the month to predict, leading to bimodal distributions for the predicted probability of being status positive. This does not happen with Stan where the latent status is represented by a continuous variable. Therefore, the predicted distributions can be different between the 2 models. This can be seen for the herd at the bottom left of Figure 9.

**Figure 8.**
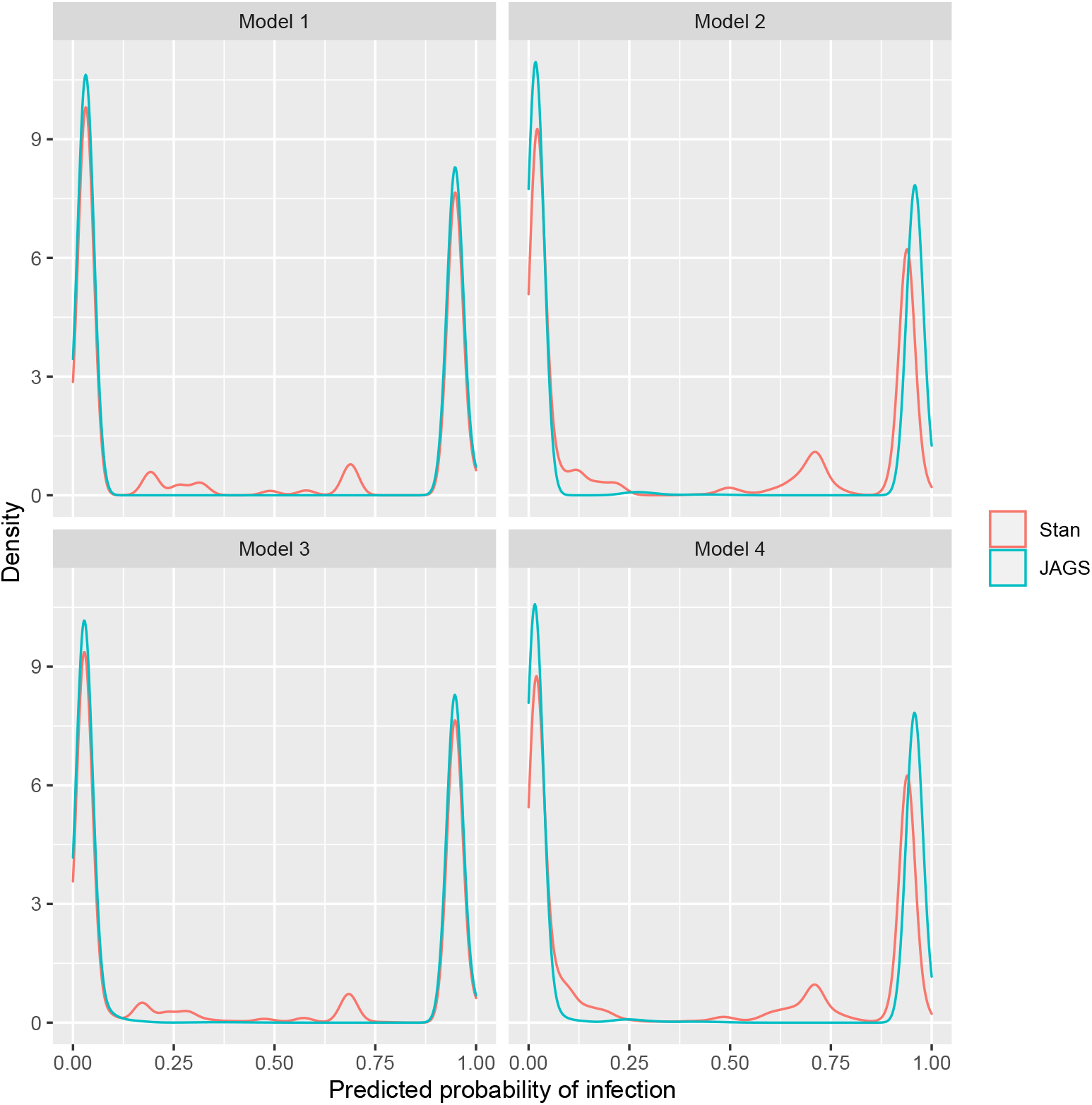
Distributions of predicted probabilities of being status positive for all herds with the 4 Bayesian models evaluated with Stan and JAGS. Model 1: Perfect test, no risk factor; Model 2: Imperfect test, no risk factor; Model 3: Perfect test, risk factor; Model 4: Imperfect test, risk factor.

**Figure 9.**
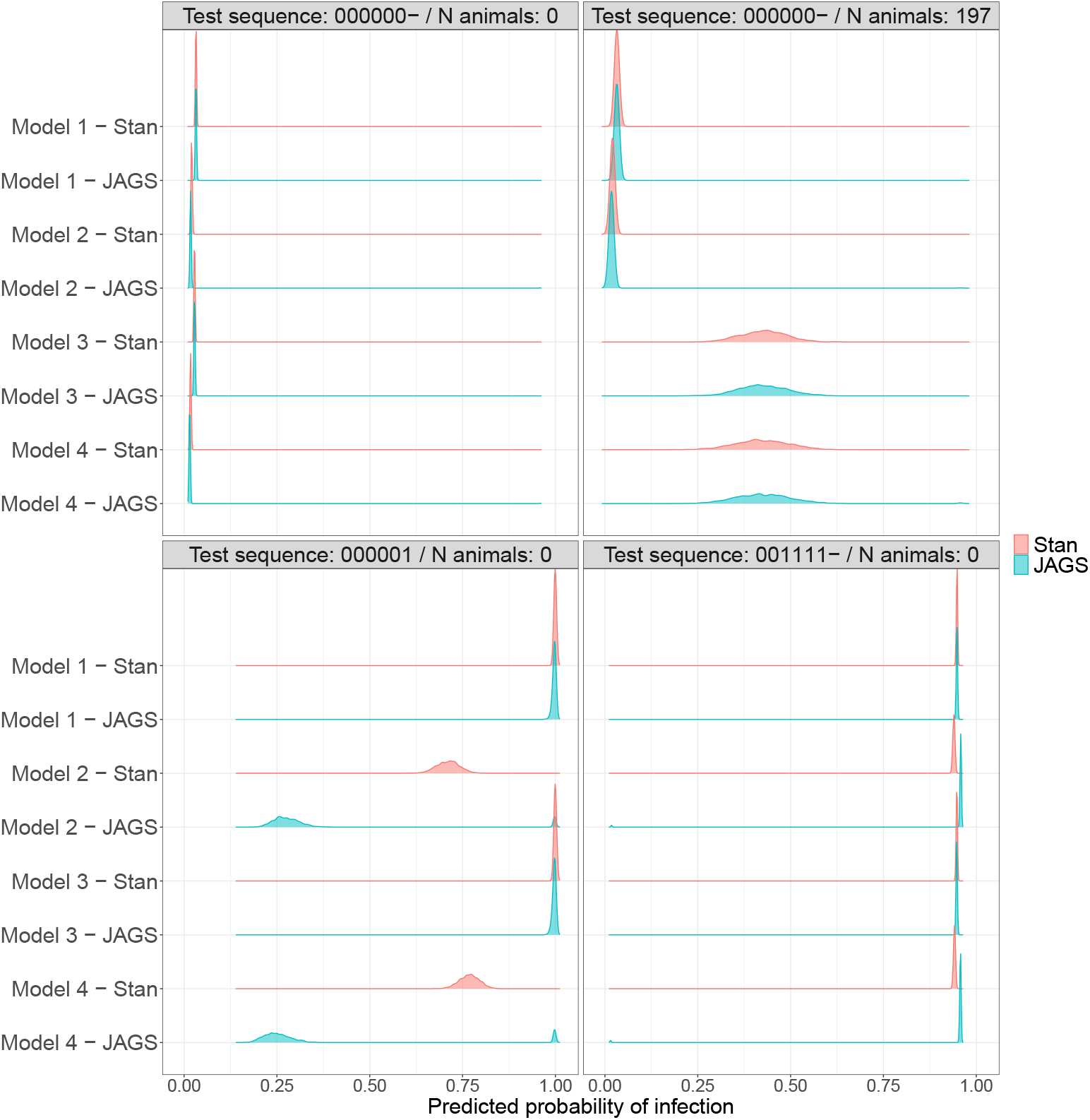
Distribution of predicted probabilities of being status positive on the month of prediction for 4 herds with the 4 models compared. Model 1: Perfect test, no risk factor; Model 2: Imperfect test, no risk factor; Model 3: Perfect test, risk factor; Model 4: Imperfect test, risk factor. The title of each panel corresponds to the sequence of test results (*-* indicates that a test result was available on the month before prediction), and the number of animals introduced 8 months before the month of prediction (risk factor).

## Discussion

This article describes a statistical framework for the prediction of an infection related status from longitudinal data generated by CPs against infectious diseases of farm animals. The statistical model developed estimates a herd level probability of being *latent status* positive on a specific month, based on input data that can vary in terms of the types of test used, frequency of testing and risk factor data. This is achieved by modelling the latent status with the same discrete time step, regardless of the frequency with which input data are available, and by modelling changes in the latent status between consecutive time steps. This model therefore fulfils one of our main objectives which was to be able to integrate heterogeneous information into the estimation. However, in order to be able to compare the output of this model run on data from different CPs, the definition of the latent status should be the same.

The model was implemented in both Stan and JAGS. The first version of the model was in JAGS, in which it was straightforward to translate the model equations into computer code. However, with this JAGS model, convergence was slow and the chains did not mix well when the prior distributions put on sensitivity and specificity were slightly wide. This led us to develop a Stan version of the model. Stan is a newer programme which uses Hamiltonian Monte Carlo for performing Bayesian inference (Carpenter et al., 2017). It was more challenging to write the model in Stan, which does not support latent discrete parameters. This was achieved by adapting a Stan implementation of the forward algorithm developed by others (Damiano et al., 2018). The Stan implementation is by comparison much faster and converges better, and should therefore be preferred.

When estimated in either JAGS or BUGS, discrete latent state models such as HMMs are known to converge slowly; and the autocorrelation in the draws from the posterior distributions is usually high. (Yackulic et al., 2020) showed that the marginalisation of the latent states considerably reduces the time needed to estimate the parameters of such models while returning the same estimates. We did not implement this approach in JAGS, although this would have been possible using the *ones trick*, as explained in the article by (Yackulic et al., 2020). The forward algorithm is a type of marginalisation that partly explains the better performance of the Stan version of the model. However, (Yackulic et al., 2020) also compared the speed of the marginalised versions of their model in different programmes and observed that Stan was orders of magnitude faster than JAGS.

In this model, the latent status is mostly defined by the prior distributions put on the different model parameters. In setting the prior distributions there are two issues: setting the distribution’s central value (mean, median …) and setting the distribution width. Using a prior distribution that does not include the true parameter value can lead to systematic error (bias) or failure of convergence. Setting prior distributions that are too wide can lead to a lack of convergence, when multiple combinations of parameter values are compatible with the data. This was a problem in the initial modelling when only the JAGS model was available. In this case, putting narrow prior distributions on test sensitivity and test specificity allowed the model to converge (results not shown). These narrow distributions imply very strong hypotheses on test characteristics.

The definition of prior distributions for test characteristics that reflect the latent status of interest is challenging (Duncan et al., 2016). This was apparent in our efforts to apply this approach to BVDV infection. For the trade of animals from herds that are free from BVDV infection, the latent status of interest was the *presence of at least one PI animal in the herd*. The test data available to estimate the probability of this event were measures of bulk tank milk antibody levels which were used to define seropositivity as a binary event. Although milk antibody level is associated with the herd prevalence of antibody positive cows (Beaudeau et al., 2001), seropositive cows can remain long after all the PIs have been removed from a herd. Furthermore, vaccination induces an antibody response which may result in vaccinated herds being positive to serological testing regardless of PI animal presence (Booth et al., 2013; Raue et al., 2011). Therefore, the specificity of BTM seropositivity, i.e. the probability for herds with no PI animals to be test negative, is less than 1. More importantly, this specificity depends on the context; i.e. on the CP. PI animals can be identified and removed more or less quickly depending on the CP, the proportion of herds vaccinating and the reasons for starting vaccination can differ between CPs. Test sensitivity can also be imperfect. Continuing with the example of bulk tank milk testing, contacts between PI animals present on the farm and the lactating herd may be infrequent, which would decrease sensitivity. In this case, the sensitivity of the testing procedure is the sensitivity of the test for the detection of seroconversion in a group of animals multiplied by the probability that the tested group has seroconverted if there is a PI animal in the herd. The probability of contact between PI animals and the lactating herd depends on how herds are organised, which could vary between CPs. This problem is alleviated when newborn calves are tested because the group of animals tested is the group in which the infectious animals are most likely to be present. Furthermore, with BTM testing, the contribution of each seropositive cow to the BTM decreases as herd size increases which can result in differences in BTM test sensitivity associated with different herd sizes between CPs.

The effects of using different prior distributions for test characteristics on latent status definition, parameter estimation and probability prediction were evaluated. In models 1 and 3, the dichotomised BTM antibody test results were modelled assuming perfect sensitivity and perfect specificity. With these assumptions, the latent status was the dichotomised test results. In Models 2 and 4, the BTM antibody test was assumed to have lower sensitivity and specificity, based on normal distributions associated with seronegativity and seropositivity identified by a mixture model. The latent status in Models 2 and 4 can therefore be described as *seropositivity*. Because overall the probability of changing status was small, assuming an imperfect sensitivity led to isolated negative test results in sequences of mostly positive test results being considered false negatives, as shown by the increase in the estimated value for *τ*_2_ between Models 1 and 2 and Models 3 and 4. This illustrates that in addition to test characteristics, status dynamics will determine the latent status within herds.

A way to obtain information on test characteristics as part of CPs could be to incorporate data from confirmatory testing into the model. In CPs, herds that test positive are usually re-tested in order to rule out a false positive test, and to identify infected animals if needed. The testing procedure used in confirmatory testing usually has a high sensitivity and a higher specificity than routine testing in relation to the gold standard. When incorporated into the model, this high quality information, in conjunction with wider prior distributions on routine testing specificity, should allow the posterior distribution of the specificity of routine testing to be revised towards the gold standard. Indeed, if a confirmatory test comes back negative, then the corresponding latent status will become negative with high probability. Given the low probability of becoming status negative between consecutive months, the latent status on the month of routine testing has an increased probability of being negative, leading to a decrease in the specificity of routine testing. Confirmatory testing data was not available for this study. We attempted to evaluate the usefulness of confirmatory testing by simulating confirmatory tests at random after an initial positive test result. The results were not convincing, because simulating test results at random was often not consistent with patterns of test results in individual herds.

Status dynamics contributed to the estimation of the latent status in several ways. Negative test results interspersed with sequences of positive test results will be classified as latent status positive (i.e. as false negatives) more often as test sensitivity decreases and *τ*_2_ increases. Positive test results interspersed with sequences of negative test results will be classified as latent status negative (i.e. as false positives) with increased frequency as test specificity and *τ*_1_ each decrease. With a perfect test (sensitivity and specificity equal to 1), the model can learn the values of *τ*_1_ and *τ*_2_ from the data, and the prior distributions put on these parameters can be minimally informative. With decreasing values for test sensitivity and specificity, the information provided through the prior distributions put on *τ*_1_ and *τ*_2_ becomes increasingly important. The informative value of *τ*_1_ and *τ*_2_ will increase as the probability of transition between latent status negative and latent status positive decrease, i.e. when *τ*_1_ is small and *τ*_2_ is high.

When data on risk factors of new infection are available, the *τ*_1_ parameter is modelled as a function of these risk factors using logistic regression. In such a case, prior distributions are put on the parameters of the logistic regression. In the application that we presented, we used a prior distribution corresponding to a low probability of new infection in the reference category (intercept: herds which introduced no animals) and we centred the prior distribution for the association with cattle introductions on a hypothesis of no association (mean = 0 on the logit scale). This allowed the model to estimate the association between the risk factor and the latent status from historical data and to use the estimated association to predict probabilities of being latent status positive on the month of prediction. The prior distributions put on test characteristics had a moderate impact on the parameter estimates. Between Model 3 and Model 4, considering an imperfect test resulted in a slightly reduced impact of the number of cattle introduced on the probability of becoming status positive (See curves at the bottom of Figure 7). The most likely explanation for this is that Model 4 allowed the highest level of discrepancy between dichotomised test result and latent status, while assuming a low probability of changing status between months. This resulted in negative test results in herds that were regularly positive to be classified as latent status positive (false negatives, associated with lower test sensitivity, see Table 3) thereby removing opportunities for new infections in herds that were regularly positive while also buying animals. This would imply that the estimated association from model 4 is more closely associated with new infections than estimates from Model 3 because herds that are regularly test positive have less weight in the estimation. It would also have been possible to base the prior distributions for the model coefficients on published literature. Unfortunately, estimates of the strengths of association between risk factors and the probability of new infection are not readily available from the published literature or are hard to compare between studies (van Roon, Mercat, et al., 2020). However, estimates from the literature could allow the prior distributions to be bounded within reasonable ranges.

The identification of the most predictive time interval between risk factor occurrence and seroconversion required the evaluation of the associations between the probability of seroconversion on a given month and risk factor occurrence over all possible intervals between this month and the 24 previous months. Although there are several Bayesian methods for such variable selection (O’Hara and Sillanpää, 2009), estimation using MCMC is time consuming and was not feasible in our case. The variables included were therefore identified with logistic models estimated by maximum likelihood for all possible lags. The approach used is related to cross-correlation maps developed for applications in ecology (Curriero et al., 2005), and similar to work conducted in veterinary epidemiology (Bronner et al., 2015). This confirmed the importance of animal introduction and neighbourhood contacts in new infections (Qi et al., 2019). However, in the Bayesian models, the 95% credibility for the association between local seroprevalence and new infection included 0 and this variable was therefore not included. The reason for this was not elucidated in this work. Other risk factors such as herd size, participation in shows or markets, the practice of common grazing have shown a consistent association with the probability of new infection by the BVDV (van Roon, Mercat, et al., 2020). These variables were not included in our model because the corresponding data were not available. One advantage of our approach is the possibility to choose candidate risk factors to include in the prediction of infection based on the data available in a given CP. The associations between the selected putative risk factors and the probability of new infection can be estimated from these data.

Given the reasonably good performance of tests for the detection of BVDV infection, the main advantage of incorporating these risk factors was not to complement the test results on a month a test was performed, but rather to enhance the timeliness of detection. Risk factors that are associated with new infection will increase the predicted probability of infection regardless of the availability of a test result. Therefore, when testing is not frequent, infected herds could be detected more quickly if risk factors of infection are recorded frequently. If the available data on risk factors of new infection captured all the possible routes of new infection, it would be possible to perform tests more frequently in herds that have a higher probability of infection as predicted by our model. In other words, our model could be used for risk-based surveillance (Cameron, 2012).

In the CP from which the current data were used, herds are tested twice a year. This could lead to a long delay between the birth of PI calves and their detection through bulk tank milk testing. We addressed this problem of *delayed detection* by proposing a method for the investigation of lagged relationships between risk factor occurrence and new infections, and by including lagged risk factor occurrences in the prediction of the probability of infection. In our dataset, herds purchasing cattle were more likely to have seroconverted 8 months after the introduction. In the Bayesian model, cattle introduction was modelled as affecting the probability of becoming status positive 8 months after the introduction. It can be argued that infection is present but not detected during this period, as the expression *delayed detection* suggests, and that the probability of infection should increase as soon as risk factor occurrence is recorded. Modelling this phenomenon would be possible by decreasing the test sensitivity for a period corresponding to the lag used in the current version of the model. This would imply that for this duration, any negative BTM test result would not provide any information about the true status regarding infection and that the herd would have an increased predicted probability of infection. This could be incorporated in future versions of the model.

There are several questions related to this modelling framework that would require further work. The model outputs are distributions of herd level probabilities of infection. Defining herds that are free from infection from these distributions will require decision rules to be developed based on distribution summaries (likely a percentile) and cut-off values. It would also be possible to model the probability of remaining infected between consecutive tests (*τ*_2_) as a function of the control measures put in place in infected herds. Another area that requires further investigations is the evaluation of the modelling framework against a simulated gold standard to determine whether it provides an added value compared to simpler methods. The availability of the model code as a Github repository allows interested users to improve or suggest improvements to our modelling framework. The model can be used to evaluate the output of disease CP thus aiding the use of output-based surveillance.

## Data accessibility

Given that the main contribution of the manuscript is of methodological nature and not about findings resulting from the analysis of the dataset, Peer Community In Animal Science has exceptionally accepted to recommend the paper without the requirement of availability of the full data set used in the manuscript. The code for the version of the modelling framework used to write the paper, as well as sample data to enable readers to use the software developed, are available here: https://doi.org/10.5281/zenodo.5292518.

## Acknowledgements

This work is part of the STOC free project that was awarded a grant by the European Food Safety Authority (EFSA) and was co-financed by public organizations in the countries participating in the study.

We thank Groupement de Défense Sanitaire de Loire-Atlantique (GDS-44) for providing the data.

Version 6 of this preprint has been peer-reviewed and recommended by Peer Community In Animal Science (https://doi.org/10.24072/pci.animsci.100007)

## Conflict of interest disclosure

The authors of this article declare that they have no financial conflict of interest with the content of this article.

Aurélien Madouasse is one of the *PCI Animal Science* recommenders.

## Footnotes

1 The *Beta*(*α* = 15, *β* = 100) distribution has a mean of 0.13 and a standard deviation of 0.03. In R, it can be plotted using the following instructions `curve(dbeta(x, 15, 100))`

2 The logit transformation is defined as 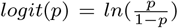 and the inverse logit transformation is defined as 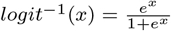. A value of 0 on the logit scale corresponds to a probability of 0.5.

3 The *logit-*^1^*Normal*(*μ* =-2, σ^2^ = 0.09) distribution has a mean of 0.12 and a standard deviation of 0.03. In R, it can be plotted using the following instructions `curve(STOCfree::dnorrn_logit(x, -2, .3))`.

4 Statuses are estimated/predicted at the herd-month level. Herd is omitted from the notation to facilitate reading. *S*_*t*_ should be read as *S*_*ht*_ where *h* represents the herd.

5 Here 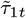 is *predicted* from herd-month specific risk factors while 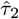is the same for all herds and *estimated* from historical data.

6 The functions used to perform this evaluation are included in the STOCfree package.

