## Supplementary file 1 for "A modelling framework for the prediction of the herd-level probability of infection from longitudinal data"

Supplementary material 1

Madouasse et al.

April 13, 2021

#### Contents

**1 Definition of a cut-off and associated charactersitics for bulk tank milk antibody testing** **2**

**2 Simplified JAGS code** **3**

**3 Stan code** **5**

### 1 Definition of a cut-off and associated charactersitics for bulk tank milk antibody testing

Test results were reported as optical density ratios (ODR). In the Loire-Atlantique CP, these ODRs are discretised into 3 categories using threshold values of 35 and 60. ODR values below 35 are associated with low antibody levels and ODR values above 60 are associated with high antibody levels. Decision regarding which herds require further testing for the identification and removal of PI animals is complex and involves the combination of test categories on 3 consecutive tests, spanning a year.

In this work, the ODR values were discretised in order to convert them into either seropositive (antibodies detected) or seronegative (no antibodies detected) outcomes. The choice of the threshold to apply for the discretisation as well as the sensitivity and specificity of this threshold for the detection of seropositivity were based on the ODR distributions from test data collected outside of the study period. Test data were available until spring 2019. However, in spring 2017, a more sensitive test was used, so this testing campaign was not analysed. Therefore, test data collected over the 4 campaigns that took place between autumn 2017 and spring 2019 were used. From these data, the overall ODR distribution was modelled as a mixture of normal distributions using the R mixdist package (Macdonald & Du, 2018). This was used to select a threshold and determine plausible sensitivity and specificity values for the selected threshold.

Test sensitivity and specificity were estimated from 5536 test results collected between 2017 and 2019. Different combinations of normal and log-normal distributions were tested to model the overall ODR distribution, using the R `mixdist` package. The best fit was obtained using 4 normal distributions (Figure 1). The 2 distributions on the left were assumed to be associated with seronegative herds and the 2 distributions on the right were assumed to be associated with seropositive herds. The cut-off of 35 used in the CP seemed to discriminate well between the distributions associated with seronegative and seropositive herds respectively, and was therefore retained in the remainder of the analysis. Using this threshold, there were 44.1% of seropositive tests between 2014 and 2016 and 40.4% between 2017 and 2019, indicating a decrease in the infection prevalence over time. With these assumptions, the test sensitivity and specificity were 0.978 and 0.948 respectively.

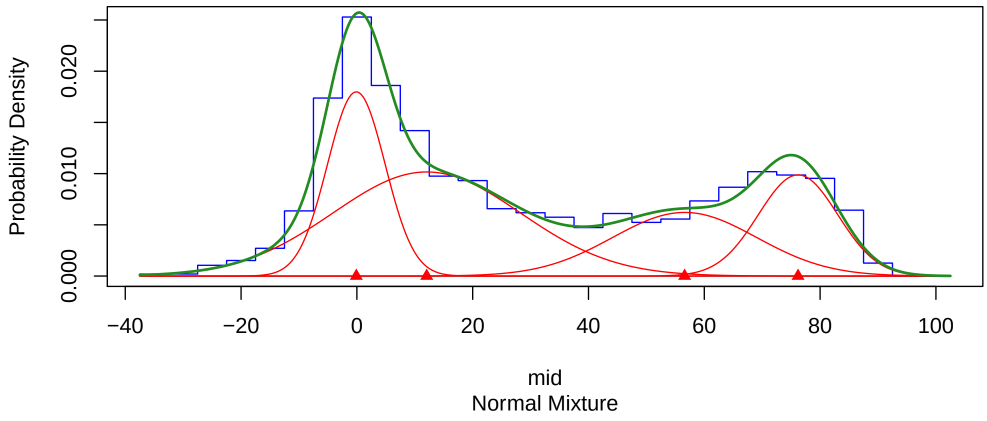


Figure 1: Distribution of the observed optical density ratios (histogram in blue) and fitted mixture of 4 normal distributions (red curves) for the bulk tank milk tests performed between 2017 and 2019. The green curve represents the overall fitted distribution.

#### 2 Simplified JAGS code

Below is simplified version of the JAGS code. This code assumes that a test result is available for each month modelled. The model implemented in the package allows for missing test results using loops to handle different possibilities. The principles are the same, but providing the actual code would make it less readable. The full model code can be generated by running the examples provided on the Github web page, by using the argument `model_code = TRUE` in the `STOCfree_JAGS()` function.

```
model{

  ## loop over all herds
  ## t1 is the vector of indices for first month of test in each herd
  ## tf is the vector of indices for last month of test in each herd
  for(i in 1:N_herds){

    ### First monthly status of each herd
    ## probability of being latent status positive for herd i at time = 1
```

```
    logit_pi1[i] ~ dnorm(logit_pi1_mean, logit_pi1_prec)

    ## latent status for herd i at time = 1
    Status[t1[i]] ~ dbern(ilogit(logit_pi1[i]))

    ## probability of being test positive given herd status
    p_test_pos[t1[i]] <- Se * Status[t1[i]] + (1 - Sp) * (1 - Status[t1[i]])

    ## test result associated with first Status => data
    test_res[t1[i]] ~ dbern(p_test_pos[t1[i]])

    ### Statuses 2 to  1 minus last
    for(t in (t1[i] + 1):(tf[i] - 1)){

      # probability of new infection
      # logistic regression
      logit(tau1[t]) <- inprod(risk_factors[t,], theta)

      ## probability of being status positive given previous status,
      ## tau1 and tau2
      pi[t] <- (1 - Status[t - 1]) * tau1[t] +
                Status[t - 1] * tau2

      ## herd status at time t
      Status[t] ~ dbern(pi[t])

      ## probability of test positive at time t
      p_test_pos[t] <- Se * Status[t] + (1 - Sp) * (1 - Status[t])

      ## test result at time t  => data
      test_res[t] ~ dbern(p_test_pos[t])

    }

    # probability of new infection
    logit(tau1[tf[i]]) <- inprod(risk_factors[tf[i],], theta)

    ## Predicted probability of infection for herd i on last month
```

```
    pi[tf[i]] <- tau1 * (1 - Status[tf[i] - 1]) +
                 tau2 * Status[tf[i] - 1]
    # probability of infection updated with test result
    predicted_proba[tf[i]] <- test_res[tf[i]] * (  # positive test result
      Se * pi[tf[i]] / (Se * pi[tf[i]] + (1 - Sp) * (1 - pi[tf[i]]))
    ) + (1 - test_res[tf[i]]) * (                  # negative test test result
        (1 - Se) * test_res[tf[i]] /
          ((1 - Se) * pi[tf[i]]  + Sp * (1 - pi[tf[i]] ) )
      )

  }

  ### Priors
  ## test characteristics
  Se ~ dbeta(Se_beta_a, Se_beta_b)
  Sp ~ dbeta(Sp_beta_a, Sp_beta_b)
  ## Status dynamics - sampling on the logit scale
  logit_tau2 ~ dnorm(logit_tau2_mean, logit_tau2_prec)
  ## logit back to the probability scale
  tau2 <- ilogit(logit_tau2)

  ## Logistic regression coefficients
  for(i_rf in 1:n_risk_factors){

    theta[i_rf] ~ dnorm(theta_norm_mean[i_rf], theta_norm_prec[i_rf])

  }

  }
```

### 3 Stan code

Stan implementation of the STOC free model. Missing test results are coded using the value 3 (instead of 0 or 1) which is handled using an if statement.

```
data{

```

```
  int<lower=1> n_herds;
  int<lower=1> herds_t1[n_herds];
  int<lower=1> herds_t2[n_herds];
  int<lower=1> herds_T[n_herds];
  int<lower=1> N;
  int<lower=0, upper=3> test_res[N];
  real<lower = 0> Se_beta_a;
  real<lower = 0> Se_beta_b;
  real<lower = 0> Sp_beta_a;
  real<lower = 0> Sp_beta_b;
  real logit_pi1_mean;
  real logit_pi1_sd;
  real logit_tau2_mean;
  real logit_tau2_sd;
  int<lower = 0> n_risk_factors;
  real theta_norm_mean[n_risk_factors];
  real theta_norm_sd[n_risk_factors];
  matrix[N, n_risk_factors] risk_factors;

}
parameters{

  real<lower = 0, upper = 1> Se;
  real<lower = 0, upper = 1> Sp;
  real<lower = 0, upper = 1> pi1;
  real<lower = 0, upper = 1> tau2;
  vector[n_risk_factors] theta;

}
transformed parameters{

  // logalpha needs to be accessible to toher blocks
  matrix[N, 2] logalpha;

  {

    // accumulator used at each time step
    real tau1[N];
```

```
  real accumulator[2];

  // logistic regression for tau1
for(n in 1:N){

tau1[n] = inv_logit(risk_factors[n,] * theta);

}


  // looping over all herds
  for(h in 1:n_herds){

    // first test in sequence
    // negative status
    logalpha[herds_t1[h], 1] = log(1 - pi1) +
                               bernoulli_lpmf(test_res[herds_t1[h]] | 1 - Sp);
    // positive status
    logalpha[herds_t1[h], 2] = log(pi1) +
                               bernoulli_lpmf(test_res[herds_t1[h]] | Se);

    // tests 2 in T in sequence
    for(t in herds_t2[h]:herds_T[h]){

      if(test_res[t] == 3){

        // transition from status negative to status negative (j = 1; i = 1)
        accumulator[1] = logalpha[t-1, 1] + log(1 - tau1[t]);
        // transition from status positive to status negative (j = 1; i = 1)
        accumulator[2] = logalpha[t-1, 2] + log(1 - tau2);

        logalpha[t, 1] = log_sum_exp(accumulator);


        // transition from status negative to status negative (j = 1; i = 1)
        accumulator[1] = logalpha[t-1, 1] + log(tau1[t]);
        // transition from status positive to status positive (j = 1; i = 1)
        accumulator[2] = logalpha[t-1, 2] + log(tau2);
```

```
          logalpha[t, 2] = log_sum_exp(accumulator);

        } else {

          // transition from status negative to status negative (j = 1; i = 1)
          accumulator[1] = logalpha[t-1, 1] +
                           log(1 - tau1[t]) +
                           bernoulli_lpmf(test_res[t] | 1 - Sp);
          // transition from status positive to status negative (j = 1; i = 1)
          accumulator[2] = logalpha[t-1, 2] +
                           log(1 - tau2) +
                           bernoulli_lpmf(test_res[t] | 1 - Sp);

          logalpha[t, 1] = log_sum_exp(accumulator);

          // transition from status negative to status negative (j = 1; i = 1)
          accumulator[1] = logalpha[t-1, 1] +
                           log(tau1[t]) +
                           bernoulli_lpmf(test_res[t] | Se);
          // transition from status positive to status positive (j = 1; i = 1)
          accumulator[2] = logalpha[t-1, 2] +
                           log(tau2) +
                           bernoulli_lpmf(test_res[t] | Se);

          logalpha[t, 2] = log_sum_exp(accumulator);

        } // if

      } // time sequence loop

    } // herd loop

  } //local

} // end of block
model{

```

```

// priors for test characteristics
     Se ~ beta(Se_beta_a, Se_beta_b);
     Sp ~ beta(Sp_beta_a, Sp_beta_b);

// priors for status dynamics
  logit(pi1) ~ normal(logit_pi1_mean, logit_pi1_sd);
  logit(tau2) ~ normal(logit_tau2_mean, logit_tau2_sd);

// priors for the logistic regression coefficients
  for(k in 1:n_risk_factors){

  theta[k] ~ normal(theta_norm_mean[k], theta_norm_sd[k]);

  }

// update based only on last logalpha of each herd
  for(i in 1:n_herds)
    target += log_sum_exp(logalpha[herds_T[i]]);

}
generated quantities{

  // variable in which predictions are stored
  real pred[n_herds];

  {
    matrix[n_herds, 2] alpha;

  // loop in which the probabilities of infection are predicted
    for(i in 1:n_herds){
      alpha[i] = softmax(logalpha[herds_T[i],]')';
      pred[i] = alpha[i, 2];
    }
  }
}
```
