## Supplementary file 2 for "A modelling framework for the prediction of the herd-level probability of infection from longitudinal data"

Supplementary material 2

Madouasse et al.

2021

### Model 1

#### Prior distributions

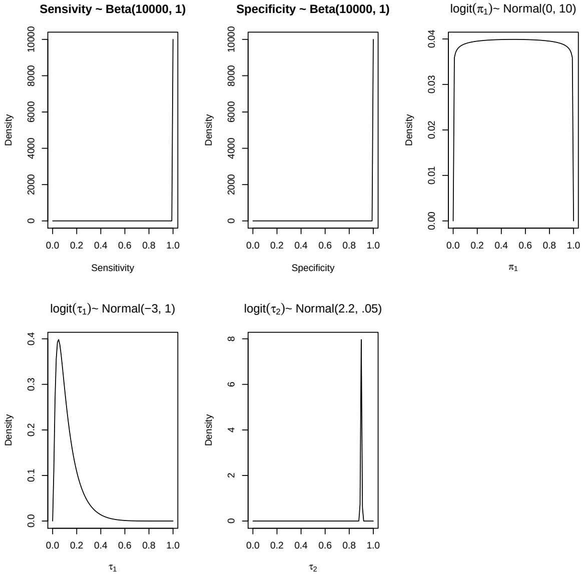

### Model 1

#### Traceplot Se - Stan

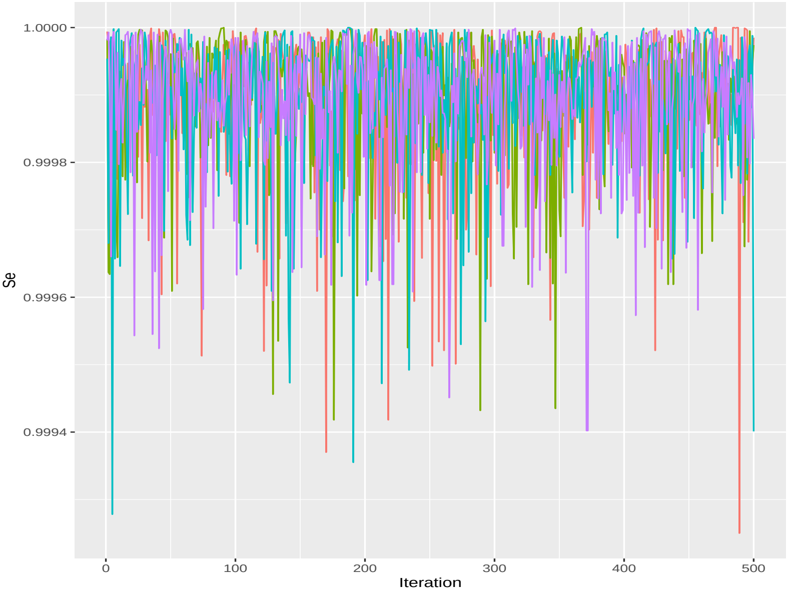

### Model 1

#### Traceplot Se - JAGS

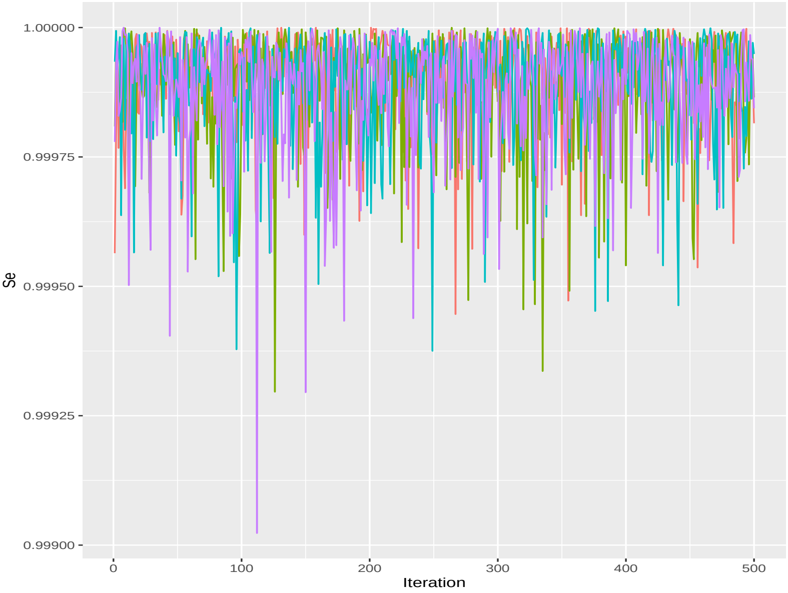

### Model 1

#### Traceplot Sp - Stan

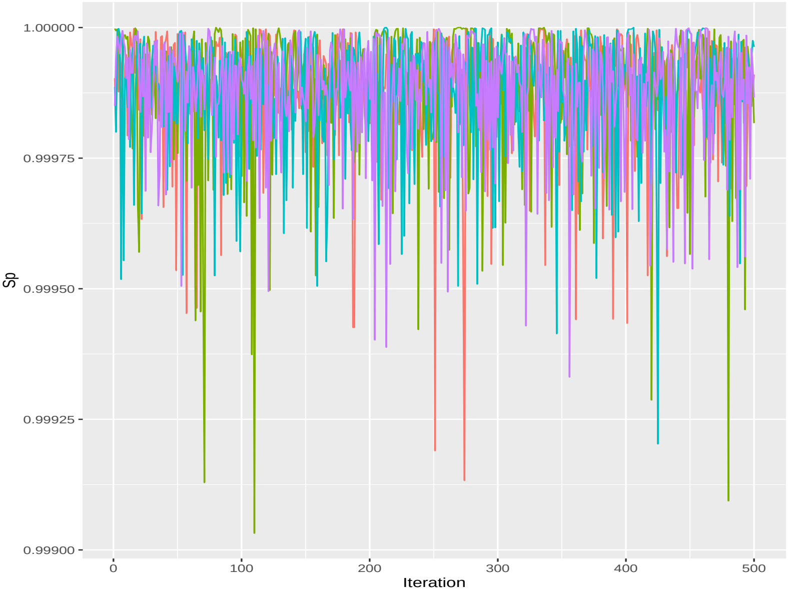

### Model 1

#### Traceplot Sp - JAGS

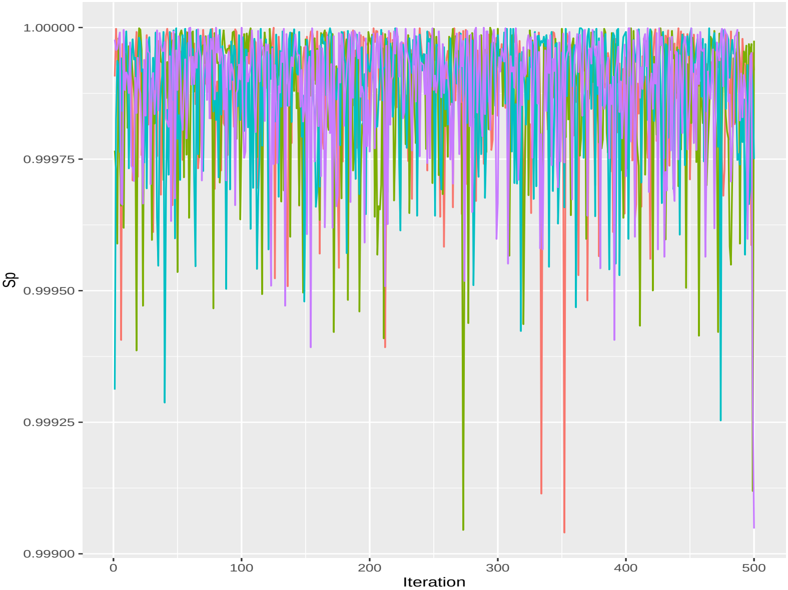

### Model 1

#### Traceplot $\tau_1$ - Stan

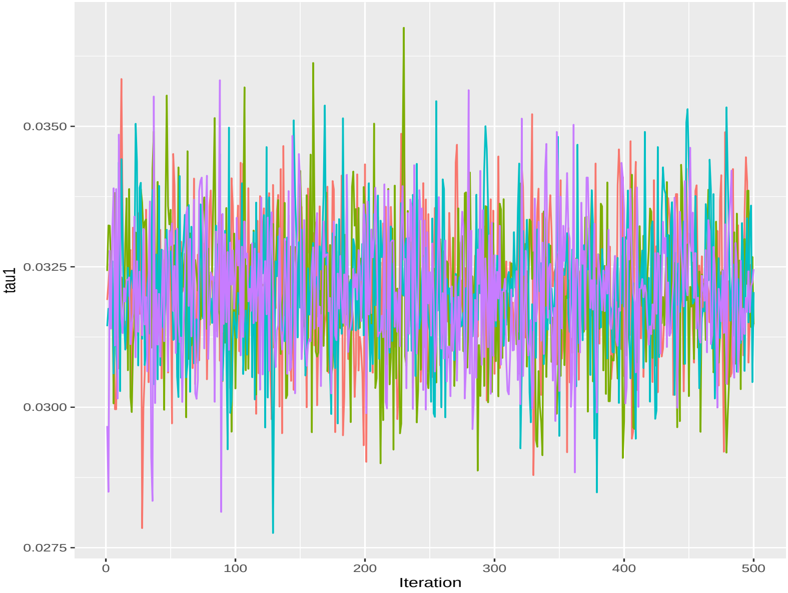

### Model 1

#### Traceplot $\tau_1$ - JAGS

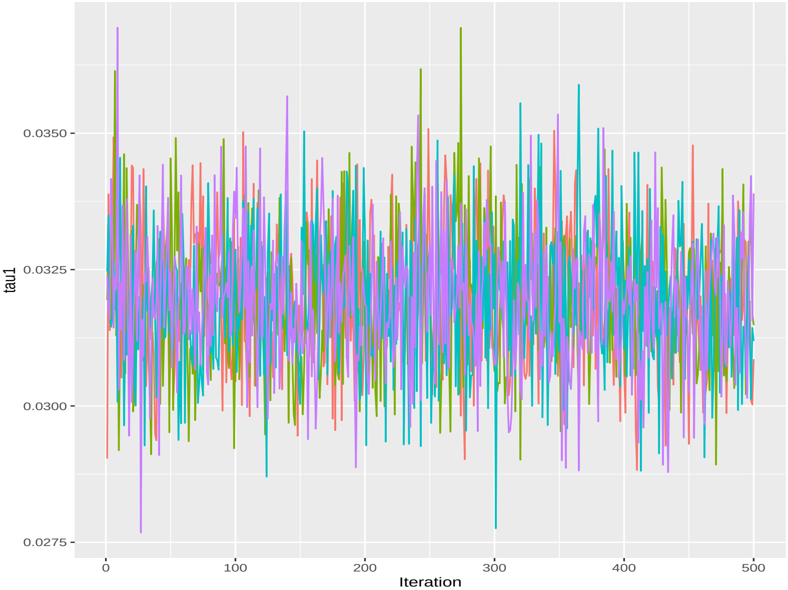

### Model 1

#### Traceplot $\tau_2$ - Stan

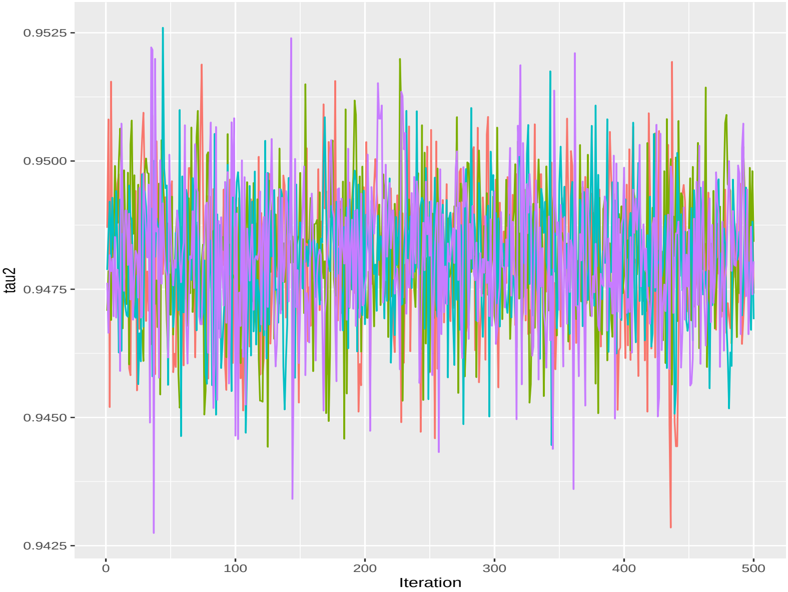

### Model 1

#### Traceplot $\tau_2$ - JAGS

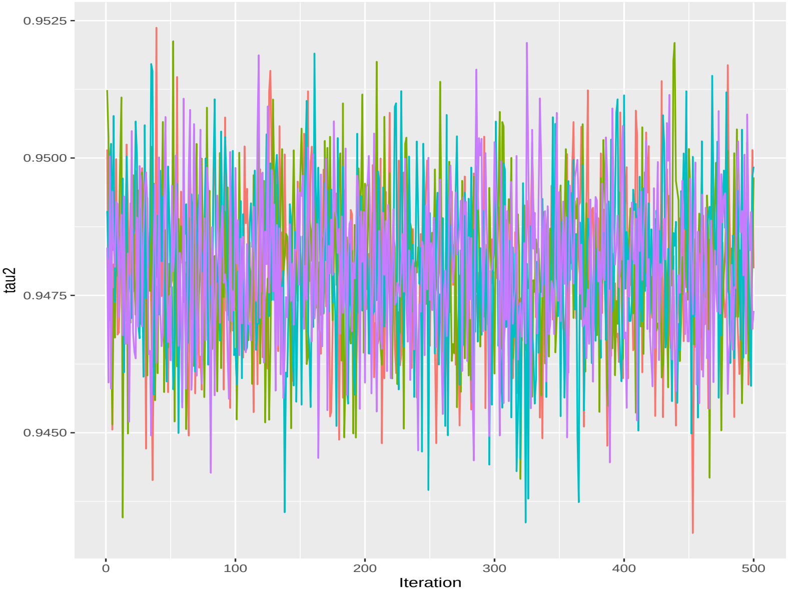

### Model 1

#### Results

##### Stan

```
##             mean           sd    median       2.5%      97.5%      ess
## Se   0.99989376 0.0001053692 0.9999270 0.99961900 0.99999700 2601.547
## Sp   0.99988042 0.0001198700 0.9999150 0.99954897 0.99999800 2723.426
## pi1  0.42245757 0.0119822438 0.4225950 0.39994568 0.44616445 2316.042
## tau1 0.03207462 0.0012292664 0.0320741 0.02975105 0.03441909 2307.151
## tau2 0.94812699 0.0013613351 0.9481435 0.94532152 0.95074420 2197.862
```

##### JAGS

```
##             mean           sd    median       2.5%      97.5%      ess
## Se   0.99988866 0.0001091851 0.9999230 0.99959282 0.99999700 1852.002
## Sp   0.99987858 0.0001248576 0.9999170 0.99954698 0.99999800 1812.019
## tau1 0.03193929 0.0012354725 0.0318992 0.02956246 0.03442477 1883.035
## tau2 0.94797153 0.0014219378 0.9479885 0.94518408 0.95078215 2000.000
```

### Model 2

#### Prior distributions

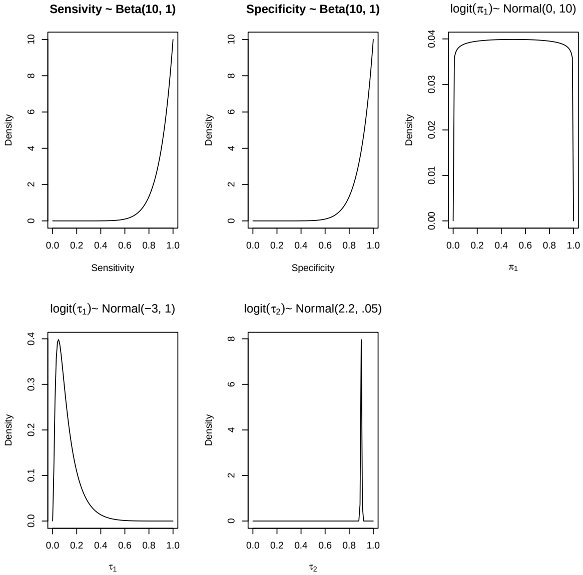

### Model 2

#### Traceplot Se - Stan

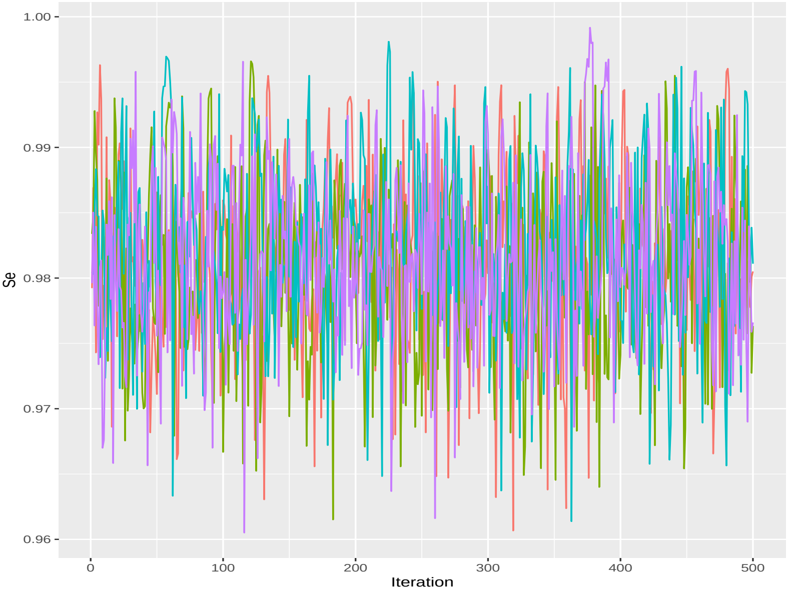

### Model 2

#### Traceplot Se - JAGS

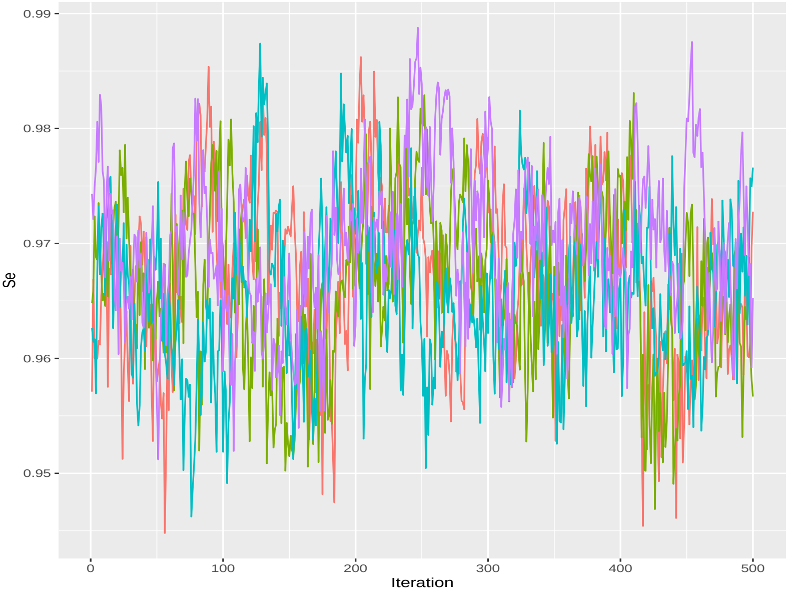

### Model 2

#### Traceplot Sp - Stan

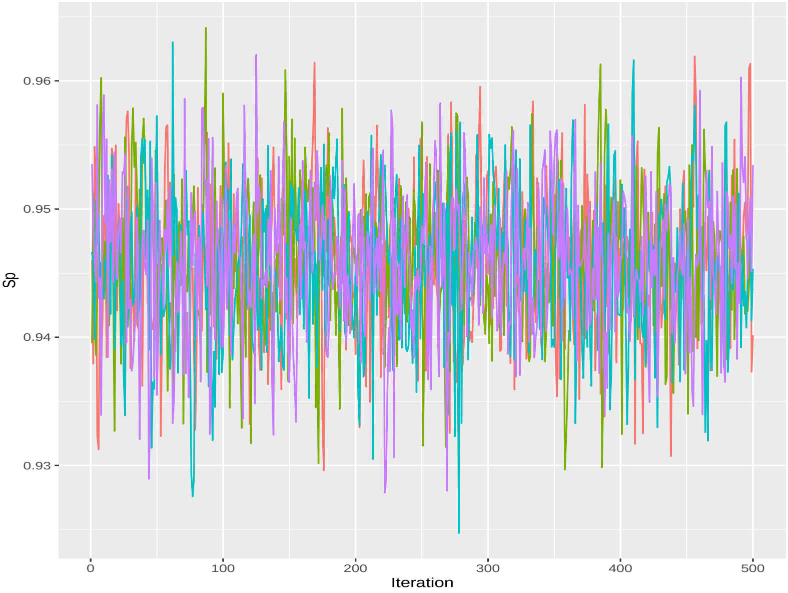

### Model 2

#### Traceplot Sp - JAGS

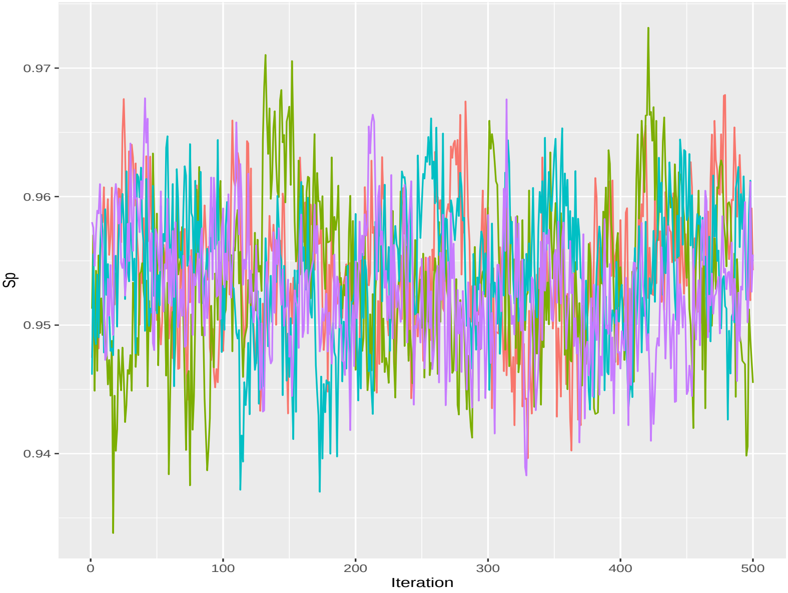

### Model 2

#### Traceplot $\tau_1$ - Stan

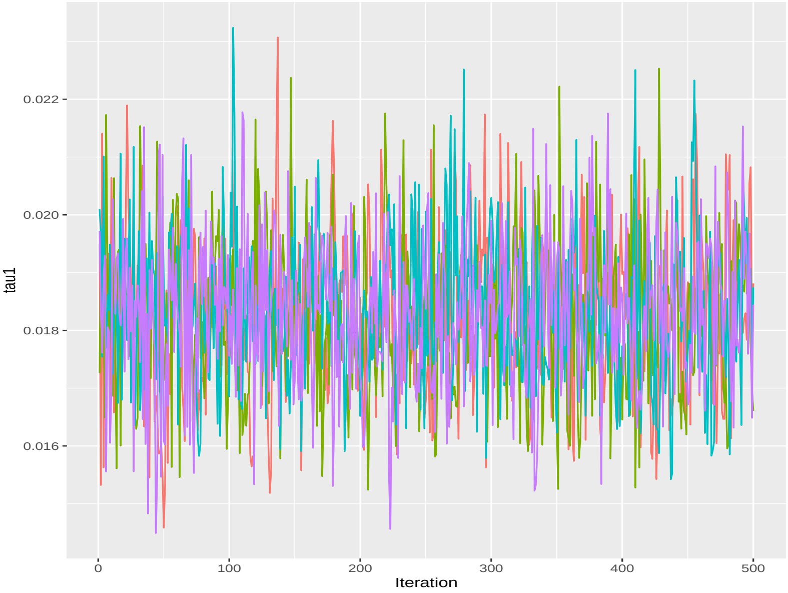

### Model 2

#### Traceplot $\tau_1$ - JAGS

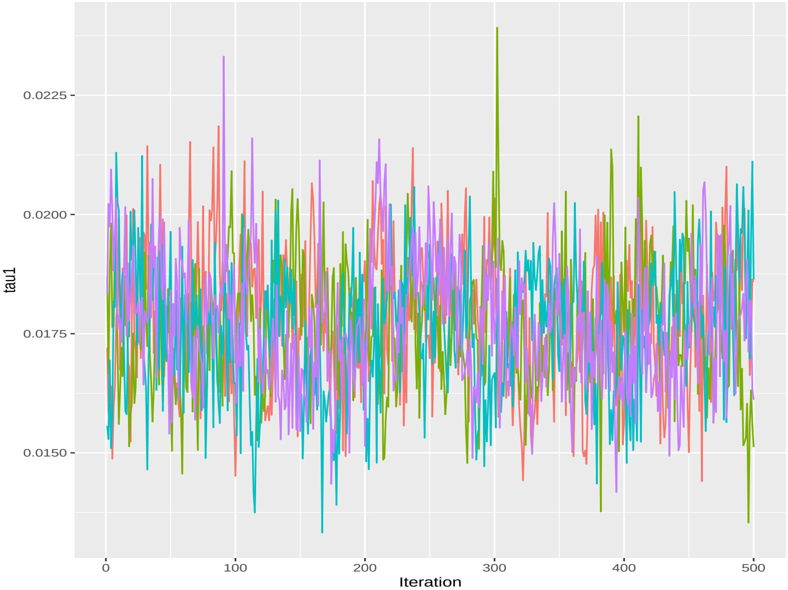

### Model 2

#### Traceplot $\tau_2$ - Stan

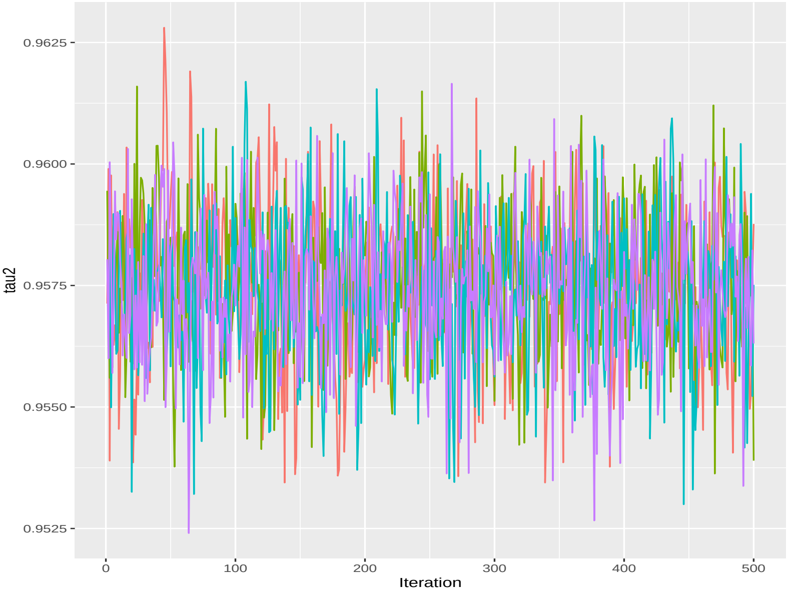

### Model 2

#### Traceplot $\tau_2$ - JAGS

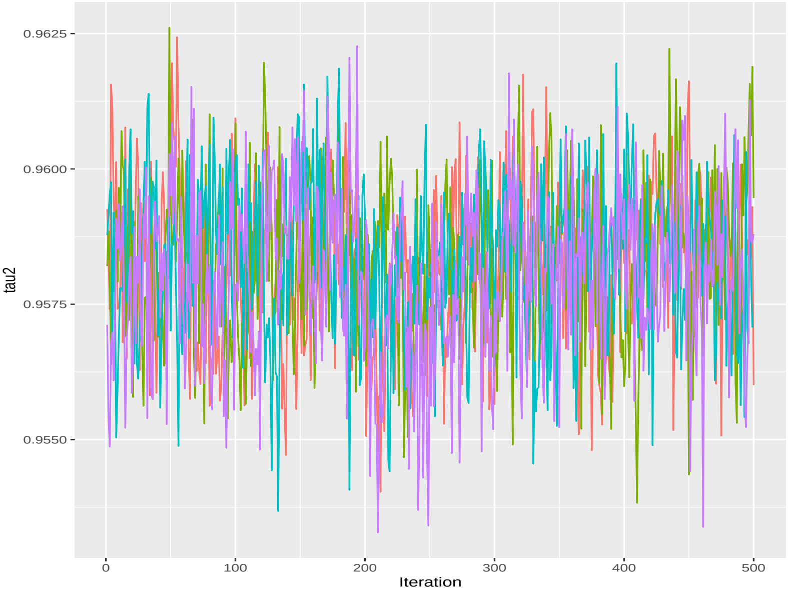

### Model 2

#### Results

##### Stan

```
##            mean          sd    median       2.5%      97.5%      ess
## Se   0.98153148 0.006772653 0.9815960 0.96764590 0.99441652 1293.904
## Sp   0.94578074 0.005760651 0.9459730 0.93387790 0.95655520 1609.149
## pi1  0.40960053 0.014337346 0.4095955 0.38182278 0.43783165 1858.271
## tau1 0.01828308 0.001325285 0.0182527 0.01582774 0.02104124 1422.944
## tau2 0.95748992 0.001502596 0.9575120 0.95447680 0.96034437 1816.834
```

##### JAGS

```
##            mean          sd     median       2.5%      97.5%      ess
## Se   0.96735005 0.007174289 0.96738050 0.95292408 0.98157328 142.2322
## Sp   0.95364573 0.005575297 0.95354450 0.94312475 0.96480888 243.2250
## tau1 0.01770141 0.001369073 0.01768865 0.01509428 0.02032662 365.3527
## tau2 0.95820726 0.001466730 0.95828350 0.95528177 0.96086675 545.3703
```

### Model 3

#### Prior distributions

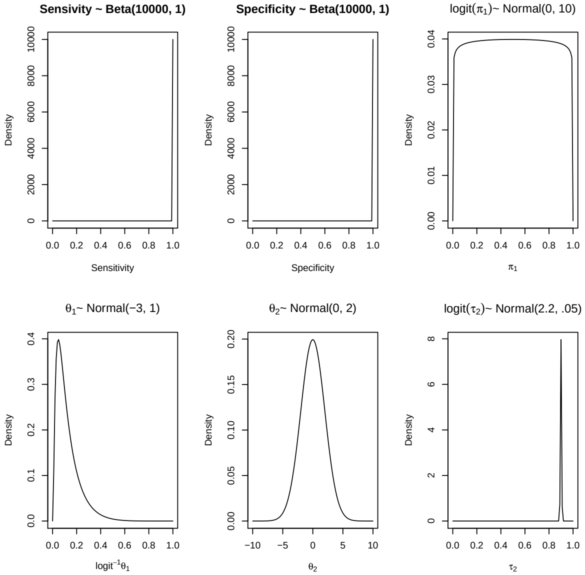

### Model 3

#### Traceplot Se - Stan

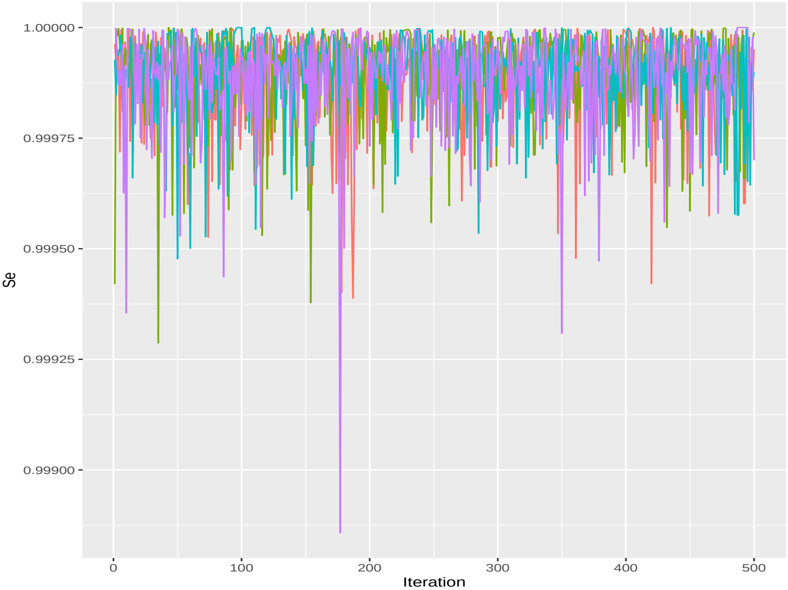

### Model 3

#### Traceplot Se - JAGS

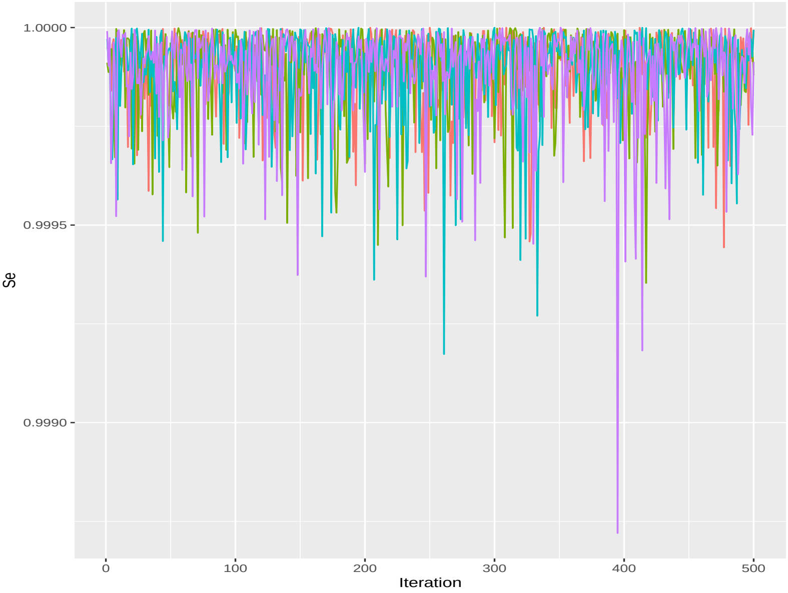

### Model 3

#### Traceplot Sp - Stan

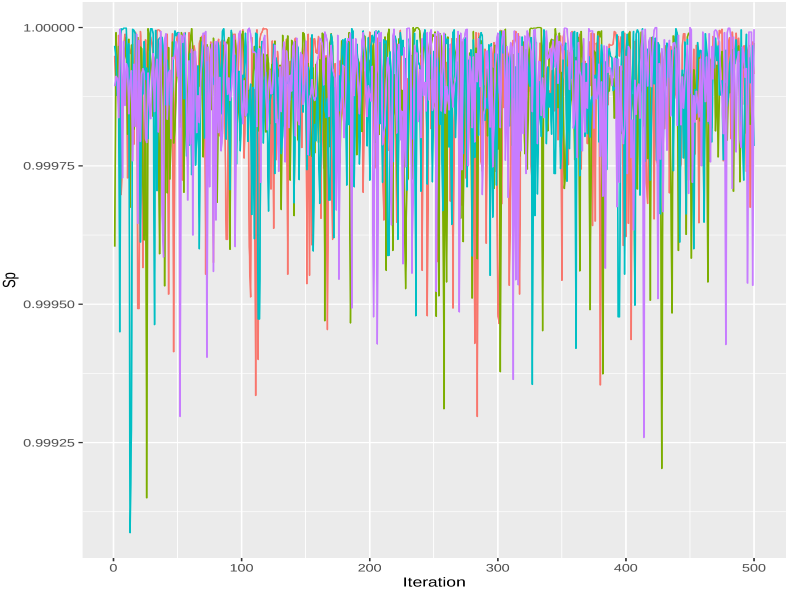

### Model 3

#### Traceplot Sp - JAGS

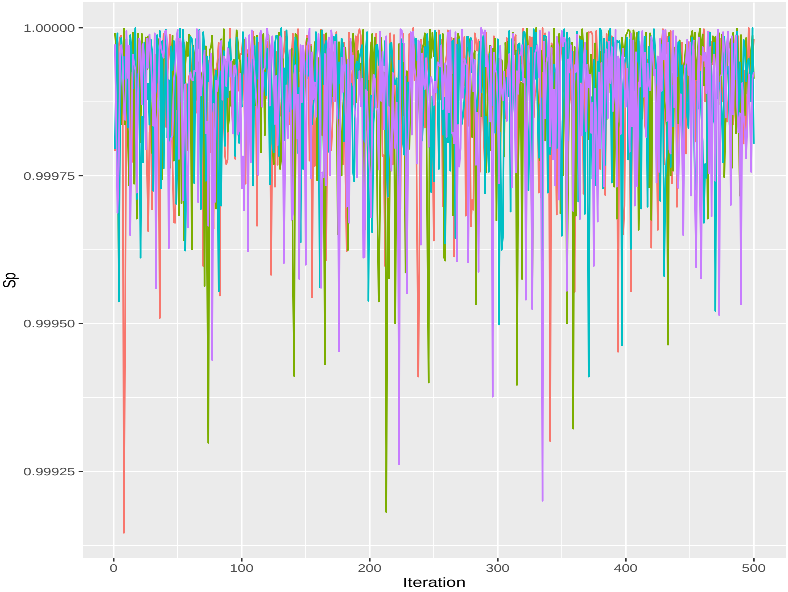

### Model 3

#### Traceplot $\theta_1$ - Stan

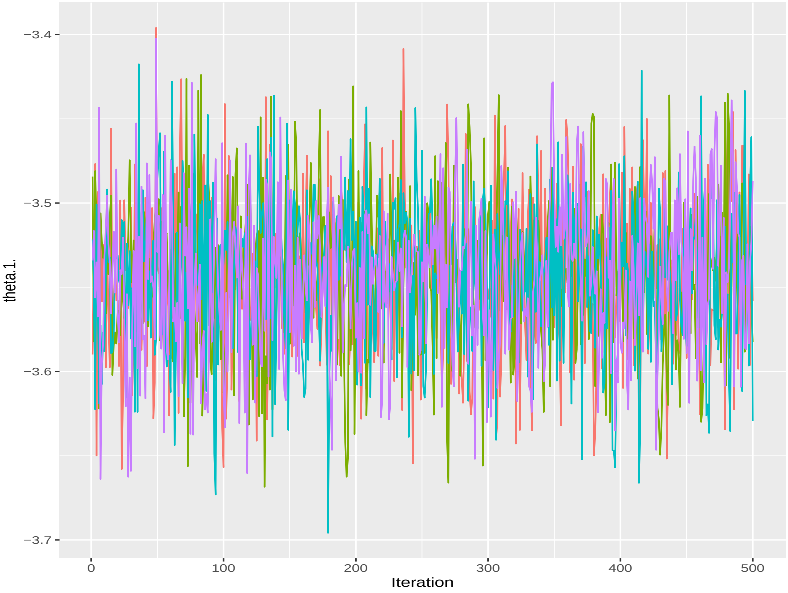

### Model 3

#### Traceplot $\theta_1$ - JAGS

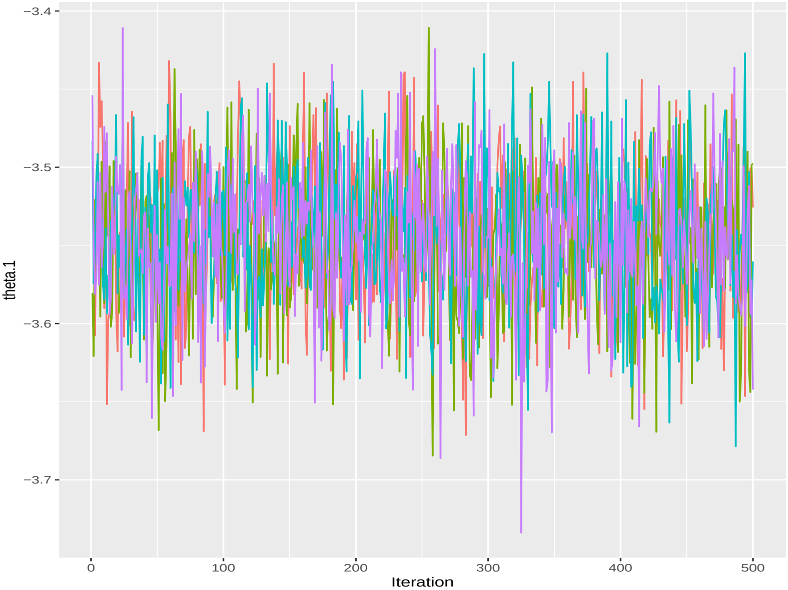

### Model 3

#### Traceplot $\theta_2$ - Stan

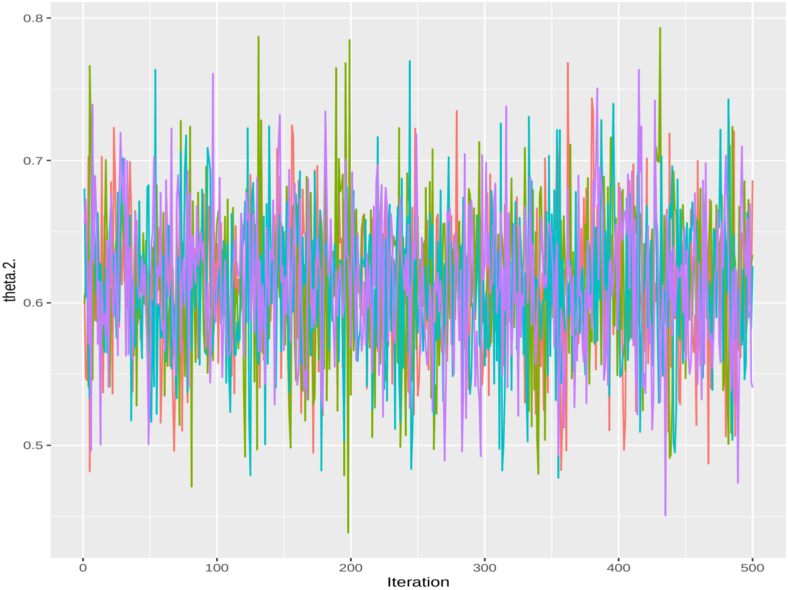

### Model 3

#### Traceplot $\theta_2$ - JAGS

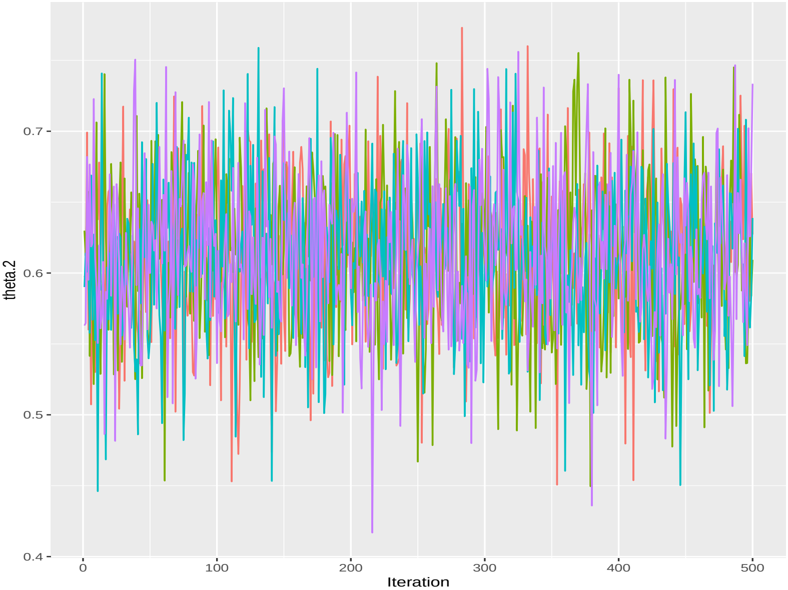

### Model 3

#### Traceplot $\tau_2$ - Stan

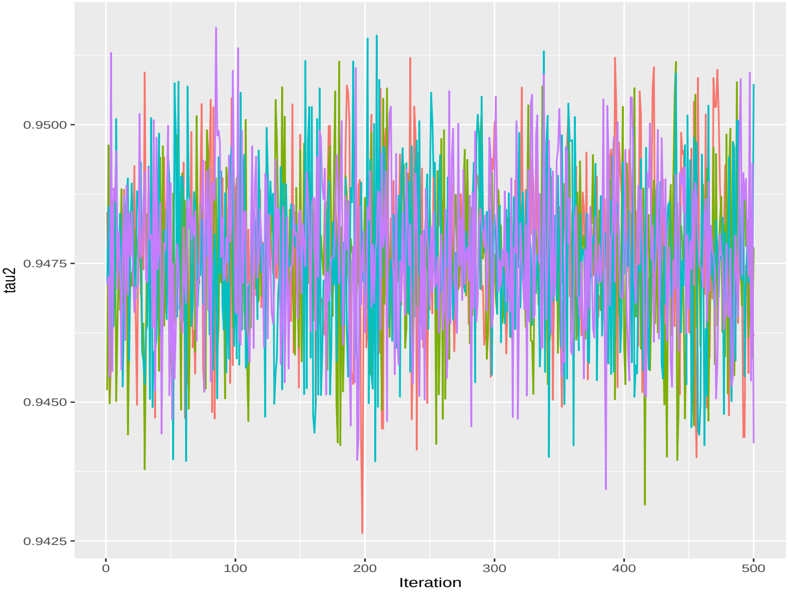

### Model 3

#### Traceplot $\tau_2$ - JAGS

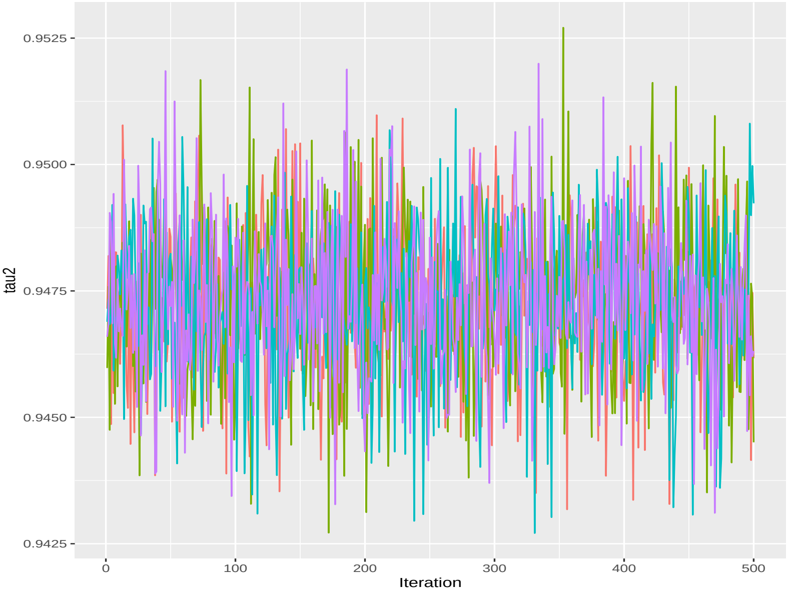

### Model 3

#### Results

##### Stan

```
##                 mean           sd     median       2.5%      97.5%      ess
## Se         0.9998923 0.0001068553  0.9999260  0.9996299  0.9999980 2541.058
## Sp         0.9998794 0.0001257076  0.9999190  0.9995129  0.9999970 2386.008
## pi1        0.4229835 0.0122642609  0.4232040  0.3980133  0.4476861 2436.371
## tau2       0.9476429 0.0014357052  0.9476455  0.9447869  0.9504940 2387.804
## theta.1. -3.5435235 0.0458758567 -3.5415950 -3.6335347 -3.4540673 1989.401
## theta.2.  0.6135751 0.0515536865  0.6153020  0.5084849  0.7162110 2566.518
```

##### JAGS

```
##                mean           sd     median       2.5%      97.5%      ess
## Se        0.9998933 0.0001127128  0.9999290  0.9995820  0.9999970 2000.000
## Sp        0.9998831 0.0001138203  0.9999170  0.9995760  0.9999960 1978.305
## theta.1 -3.5452866 0.0452831075 -3.5447350 -3.6363005 -3.4575155 2026.991
## theta.2  0.6129062 0.0539827551  0.6127870  0.5079346  0.7207263 1735.106
## tau2     0.9472999 0.0014943146  0.9473445  0.9442210  0.9502642 2114.463
```

### Model 4

#### Prior distributions

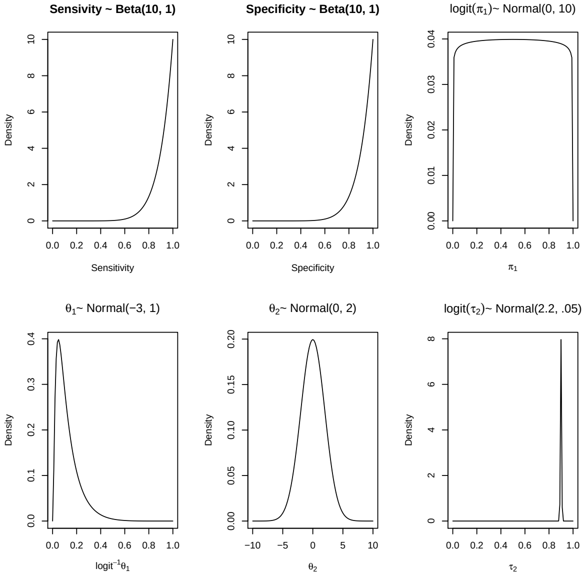

### Model 4

#### Traceplot Se - Stan

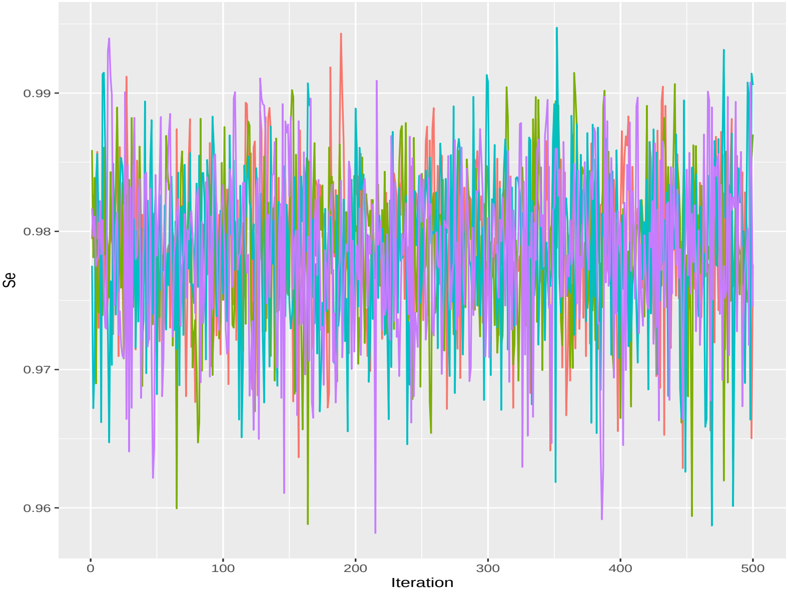

### Model 4

#### Traceplot Se - JAGS

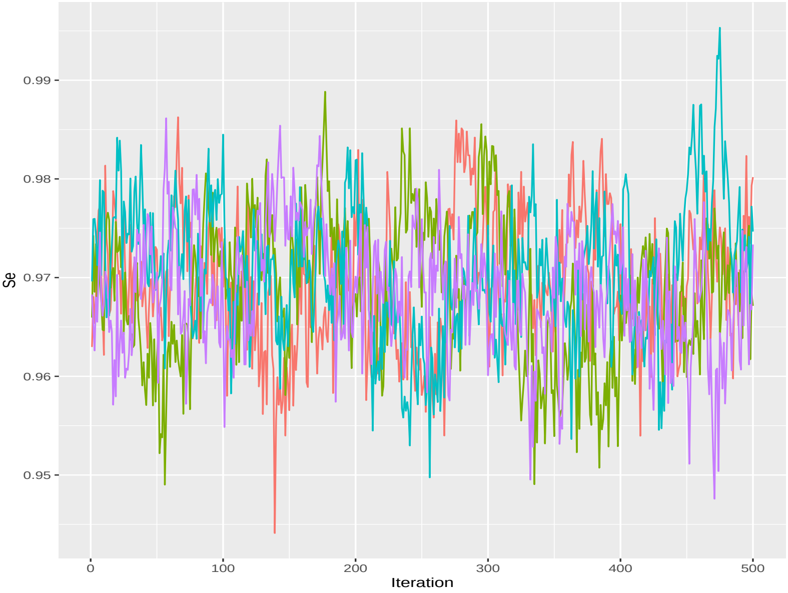

### Model 4

#### Traceplot Sp - Stan

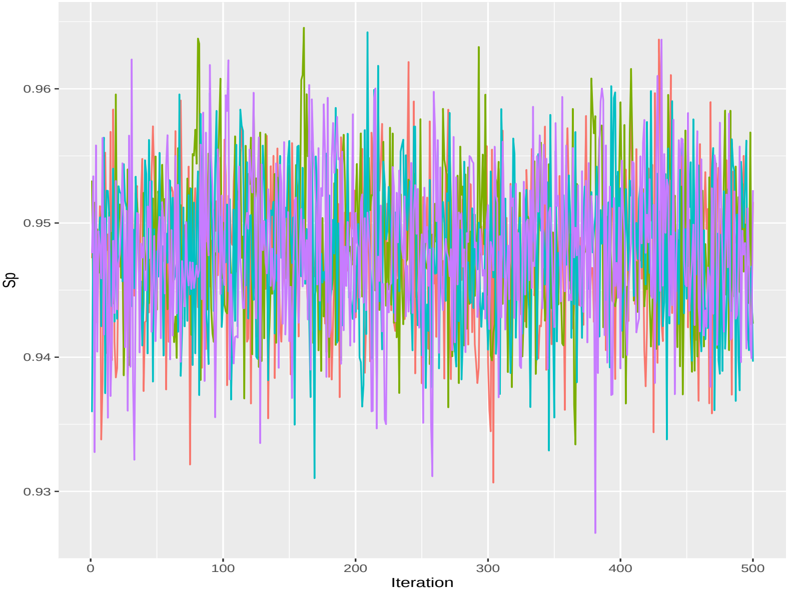

### Model 4

#### Traceplot Sp - JAGS

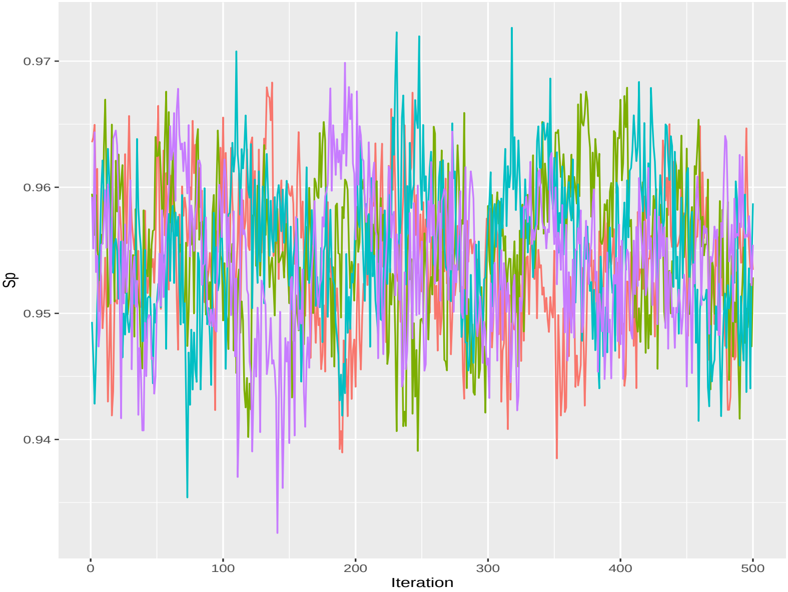

### Model 4

#### Traceplot $\theta_1$ - Stan

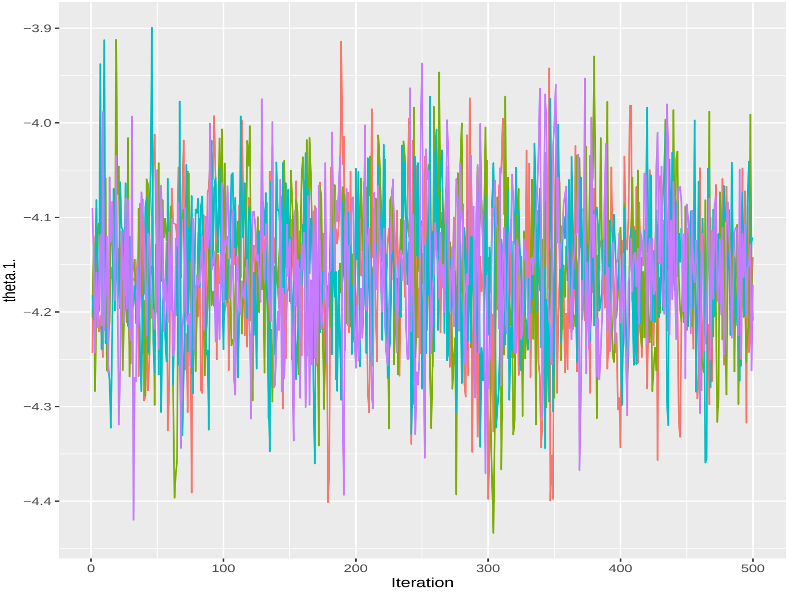

### Model 4

#### Traceplot $\theta_1$ - JAGS

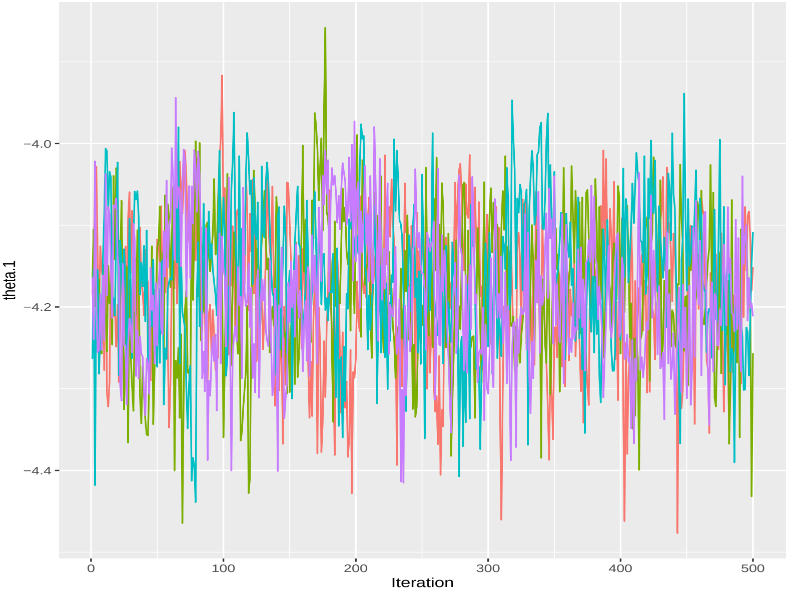

### Model 4

#### Traceplot $\theta_2$ - Stan

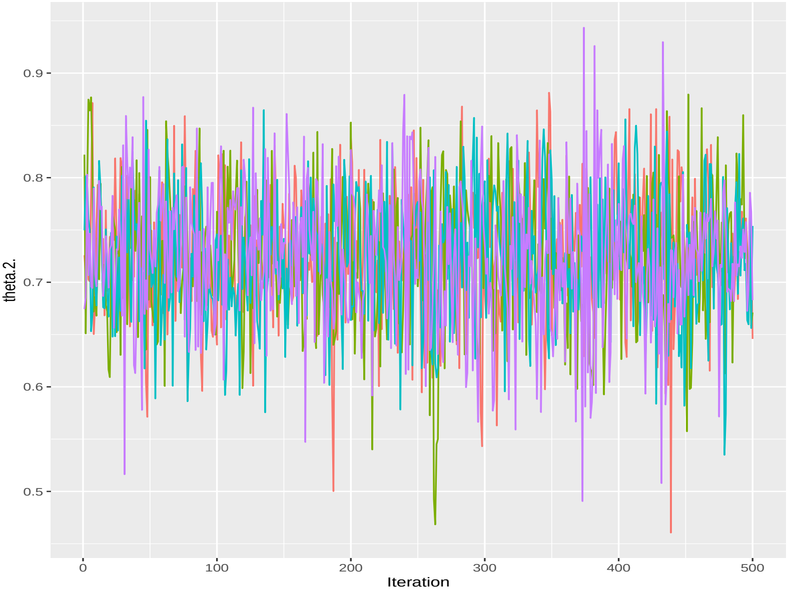

### Model 4

#### Traceplot $\theta_2$ - JAGS

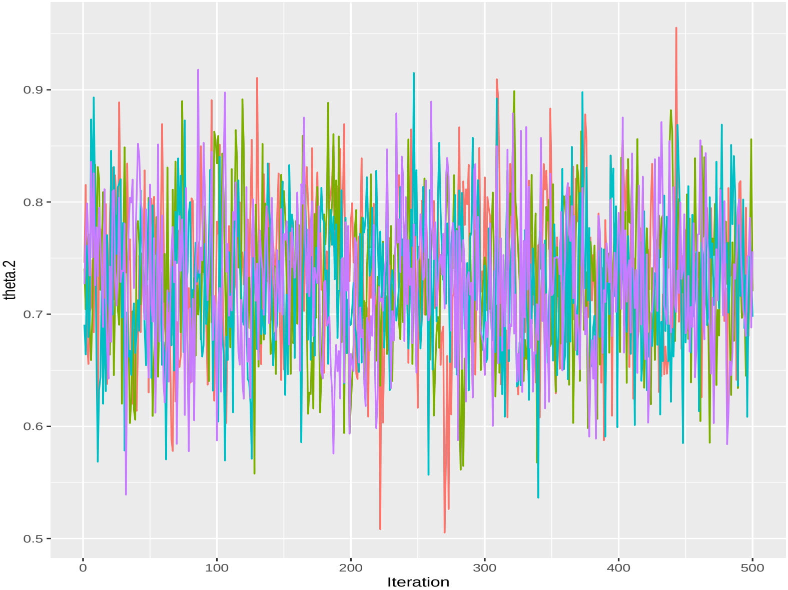

### Model 4

#### Traceplot $\tau_2$ - Stan

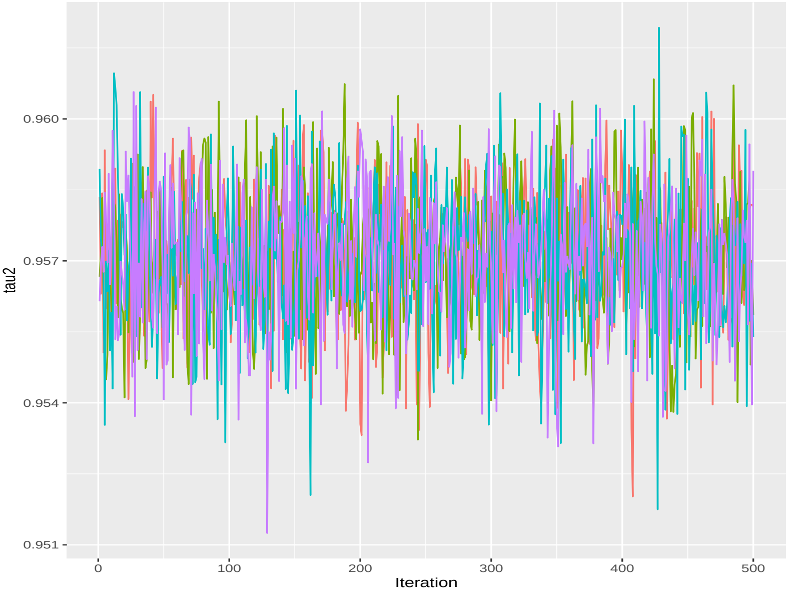

### Model 4

#### Traceplot $\tau_2$ - JAGS

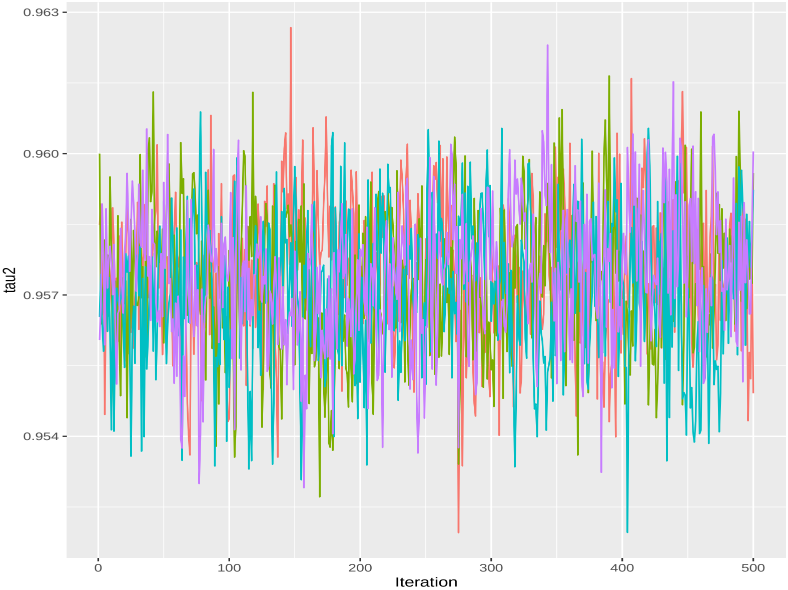

### Model 4

#### Results

##### Stan

```
##                  mean          sd     median       2.5%      97.5%      ess
## Se          0.9787592 0.005700809  0.9790445  0.9664178  0.9892195 2087.078
## Sp          0.9480368 0.005590873  0.9479030  0.9371573  0.9590133 2007.011
## pi1         0.4107576 0.013008345  0.4108420  0.3843662  0.4364940 2173.017
## tau2        0.9570797 0.001471666  0.9571275  0.9540979  0.9598452 3039.028
## theta.1. -4.1586092 0.078906029 -4.1559800 -4.3198643 -4.0032375 1432.438
## theta.2.  0.7244467 0.062941083  0.7246880  0.5958777  0.8424609 1802.306
```

##### JAGS

```
##                mean          sd     median       2.5%      97.5%       ess
## Se        0.9693652 0.006778411  0.9693805  0.9563966  0.9824612  145.0035
## Sp        0.9546281 0.005765281  0.9547860  0.9430777  0.9651936  174.6555
## theta.1 -4.1771472 0.088496446 -4.1773050 -4.3575447 -4.0141232  404.9377
## theta.2  0.7309639 0.064165430  0.7313855  0.6064481  0.8555339 1057.3814
## tau2      0.9573082 0.001506189  0.9573440  0.9541348  0.9601620  566.9616
```

### Predicted probabilities of status positive

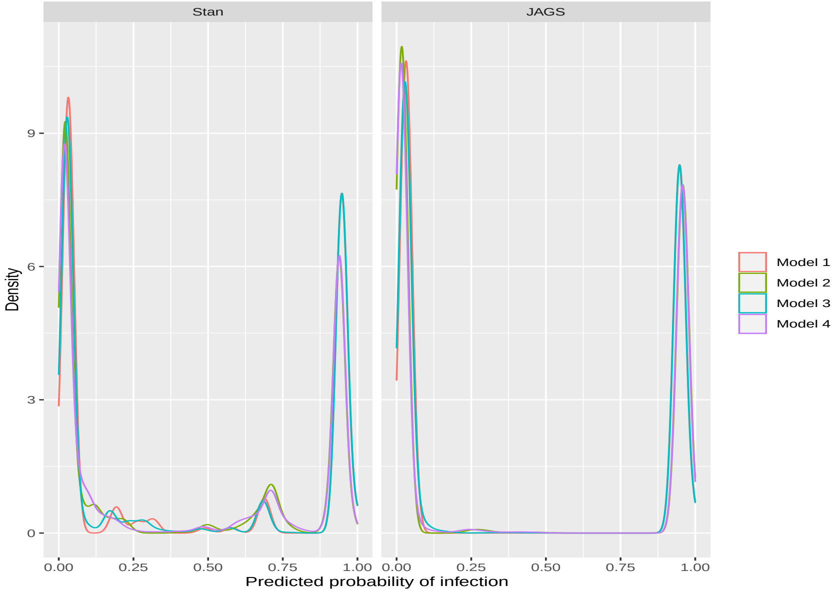
